# Natural and breeding selection converge on overlapping haplotypes with divergent directions and outcomes in wheat

**DOI:** 10.64898/2026.03.28.714077

**Authors:** Xiaoming Wang, Jesus Quiroz-Chavez, Ricardo H Ramírez-González, Zile Xiong, Xue Shi, Shengbao Xu, Sacha Przewieslik Allen, Shifeng Cheng, Nikolai Adamski, Cristobal Uauy

## Abstract

Understanding how natural and breeding selection interact to shape crop genomes is essential for improving resilience under climate change. Here, we applied a *k*-mer-based, alignment-free haplotype assignment approach to whole-genome resequencing data from 827 wheat landraces representing seven geographic groups and 208 modern cultivars. We identified haplotypes associated with local adaptation that were enriched in specific agroecological regions, many of which were derived from wild-relative introgressions. Comparative analyses revealed that natural and breeding selection largely targeted overlapping haplotype sets, but often drove them in opposite directions. Notably, haplotypes conferring adaptive advantages were frequently associated with negative regulation of agronomic traits, explaining their reduced prevalence among breeding-selected haplotypes. These results reveal the genomic basis of trade-offs between environmental adaptation and productivity and offer a framework for exploiting adaptive diversity in wheat improvement.

## Introduction

The global spread of crop species from their centres of origin represents one of the most remarkable events in agricultural history. During this migration, crops encountered diverse geographic environments, and their genomes underwent profound changes to enable adaptation to new ecological niches^1–5^. While substantial progress has been made in understanding the genomic basis of crop domestication and adaptation, critical questions remain unresolved regarding how local adaptation and subsequent selective pressures are reflected in haplotype diversity across the genome^6–8^. In particular, the identification of adaptive haplotypes has the potential to provide indispensable genetic resources for addressing the challenges posed by climate change^7,9,10^.

Beyond natural adaptation, crop genomes have also been shaped by human-mediated breeding. Modern breeding programs have accelerated the fixation of alleles conferring desirable agricultural traits, often narrowing allelic diversity compared with landraces^11–13^. One recent study has revealed that approximately 31% of haplotypes are influenced by both natural and breeding selection, suggesting an interplay between these two forces^9^. Due to the difference in objectives, natural selection drives local adaptation, whereas breeding selection prioritizes yield and quality improvement; the trajectories of these two evolutionary forces may diverge. This interplay creates both tension and opportunity: reconciling adaptive variation preserved by natural selection with the goals of modern breeding will be critical for achieving future crop resilience^10^.

Wheat provides an exceptional system for investigating these dynamics. Originating in the Fertile Crescent, hexaploid bread wheat (*Triticum aestivum* L.) emerged around 10,000 years ago through hybridization events involving multiple wild progenitors, making it a relatively young crop^3,9,14^. Its subsequent global spread exposed it to diverse agroecological environments, requiring rapid genomic responses ^3,6,9,15^. In parallel, modern breeding has imposed strong artificial selection pressures, particularly during the Green Revolution and subsequent international improvement programs^13,16,17^. Advances in large-scale resequencing and pan-genomics now offer unprecedented opportunities to resolve how these evolutionary forces have shaped wheat genomes^17–20^.

In this study, we leveraged a large-scale resequencing dataset comprising 827 landraces and 208 modern cultivars from diverse geographic regions^13^. Using a *k*-mer-based haplotype detection method independent of sequence alignment, we identified haplotypes associated with local environmental adaptation and examined their trajectories under breeding selection. Our analyses reveal that although natural and breeding selection often target the same haplotypes, they frequently act in opposite directions, largely because adaptive haplotypes tend to negatively affect agronomic performance. We also demonstrate that natural introgressions from wild relatives substantially contributed to wheat’s adaptive landscape. Together, these findings provide new insights into the genomic basis of wheat’s global spread, highlight the evolutionary tension between natural and breeding selection, and underscore the potential of haplotype diversity for breeding climate-resilient cultivars.

## Results

### Long-range haplotypes reveal a high-resolution presence/absence map between wheat landraces and modern cultivars

To comprehensively assess the genetic diversity of Watkins landraces, we reanalyzed the genetic variation using the independent *k*-mer-based approach developed in our previous studies^13,21,22^. Cleaned sequencing reads from 827 landraces (LAs) and 208 modern cultivars (MCs), together with the wheat reference genome, were used to construct 31-bp *k*-mer databases, respectively. Pairwise comparisons between sequencing and reference sequence *k*-mers generated Identity-by-State Python (IBSpy) values across 50-kb non-overlapping windows, resulting in a variation matrix for all 1,035 accessions^23^. Principal component analysis (PCA) separated the seven LA groups and placed MCs closest to AG2 and AG5, consistent with results from conventional SNP-based methods^13^ (Fig. 1a; Supplementary Fig. 1a). Haplotypes were assigned by clustering 20 consecutive IBSpy values using affinity propagation^13^, producing a genome-wide haplotype matrix at 1-Mb resolution^23^. As expected, haplotype diversity was much higher in LAs than in MCs, particularly in distal chromosomal regions (Fig. 1b; Supplementary Fig. 1b).

**Fig. 1:**
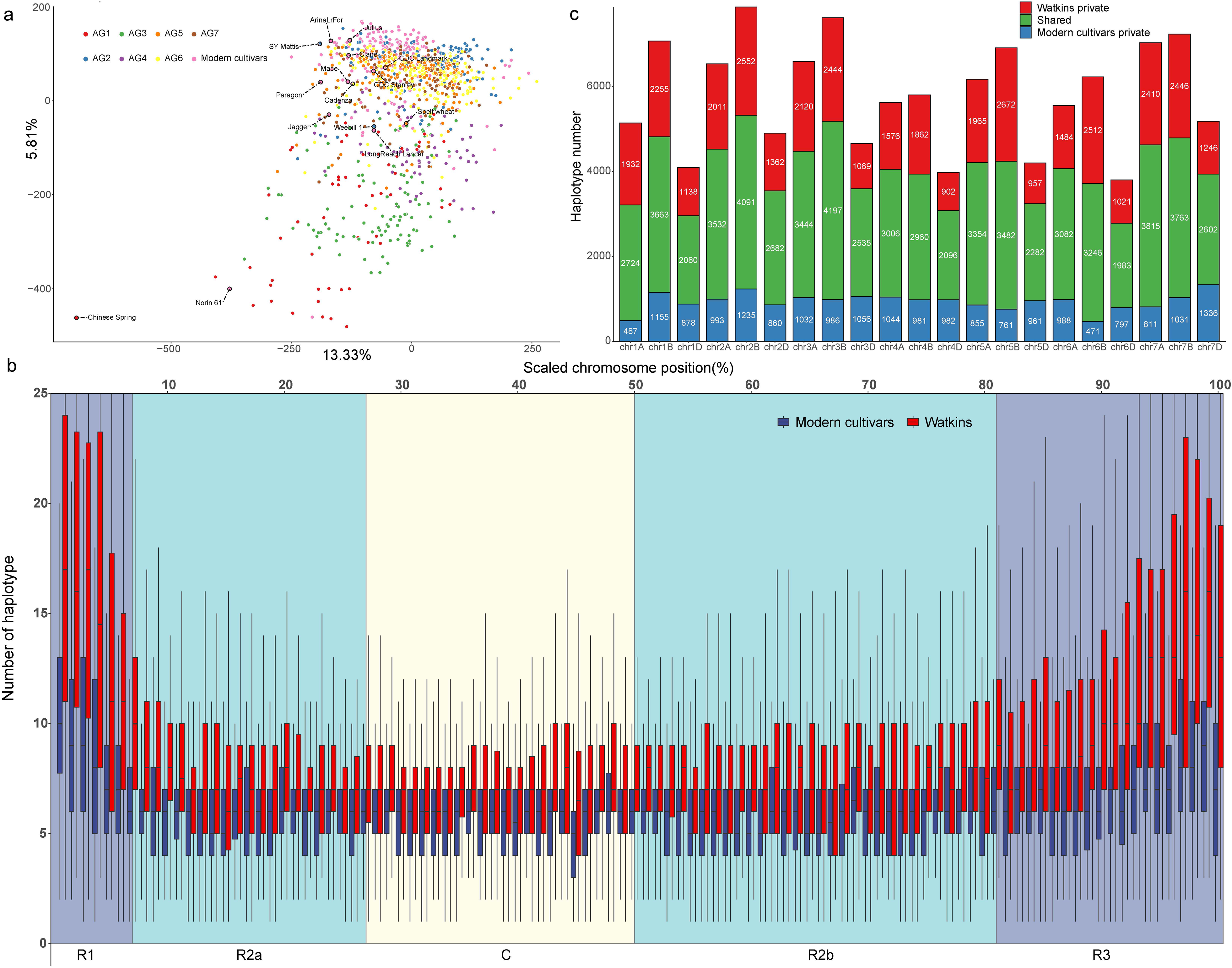
Identification of landrace- and modern cultivar-private haplotypes using *k*-mer-based IBSpy variations and haplotypes with Chinese Spring as reference. a, PCA (Principal Component Analysis) plot of the first two components calculated using the genome-wide IBSpy variations of 827 landraces and 244 modern cultivars. The seven groups of Watkins and modern cultivars were defined in our previous study^13^, and the 15 previously described wheat cultivars from the 10+ Wheat Genomes Project^17^ were highlighted. **b**, Distribution of haplotype number per Mb across all 21 chromosomes (positions scaled to % of maximum chromosome length). Boxplots show distributions of 1% bins. The four chromosomal compartments were defined based on the recombinations^18^. **c**, Number of landraces (Watkins)- and modern cultivars (Modern)-private haplotypes and the shared haplotypes between these two groups across all 21 chromosomes.

This high-resolution haplotype framework allowed precise detection of genomic regions that were lost or gained during wheat breeding. We identified 37,936 LA-private haplotypes (present in ≥3 LAs but ≤1 MC) and 19,700 MC-private haplotypes (present in ≥2 MCs but ≤2 LAs), accounting for 37.0% and 23.4% of their major haplotypes, respectively (Supplementary Tables S1, S2). Subgenome B contained the largest proportion of LA-specific haplotypes (44.1%; Fig. 1c, Supplementary Fig. 1c), consistent with its higher genetic diversity^13,16,24^. Most private haplotypes were located in distal regions (Supplementary Fig. 2). Projecting the private haplotypes to the chromosomes of each accession revealed large private haplotype blocks in both LAs and MCs (Supplementary Fig. 3), including the well-characterised human-derived introgressions such as 1RS·1BL from *Secale cereale* in 28 MCs^25–27^ (Supplementary Fig. 4), and the 2NvS fragment from *Aegilops ventricosa* in 11 MCs^17^ (Supplementary Fig. 5). Together, these analyses establish a genome-wide presence/absence haplotype map at 1-Mb resolution, highlighting both untapped diversity in LAs and introgression segments in MCs.

### Identification of local adaptation haplotypes during wheat expansion

Between 2000 and 500 BCE, wheat expanded from its origin in the Fertile Crescent to diverse environments, shaped by both natural and human selection, and developed a variety of adaptive traits that enabled it to thrive in new environments^6,28,29^. To uncover genomic regions associated with this expansion, we developed a haplotype frequency differential-based method to identify local adaptation-related haplotypes (LAr-Haps), defined as significantly higher-frequency haplotype in a given geographic group (AG1-AG7)^13^ relative to others (Fig. 2a; Methods). We identified LAr-Haps in each of the seven regions based on 21 pairwise comparisons (Fig. 2b). AG4 and AG1, representing the Middle East and East Asia, contained the most LAr-Haps (11,122 and 10,511), followed by AG3 (9,233), AG2 (8,396), AG7 (7,659), AG6 (6,662), and AG5 (5,133) (Supplementary Table S3 and Fig. 6). Geographically neighbouring regions shared more LAr-Haps, such as AG4, AG1 and AG3, consistent with local environmental adaptation.

**Fig. 2:**
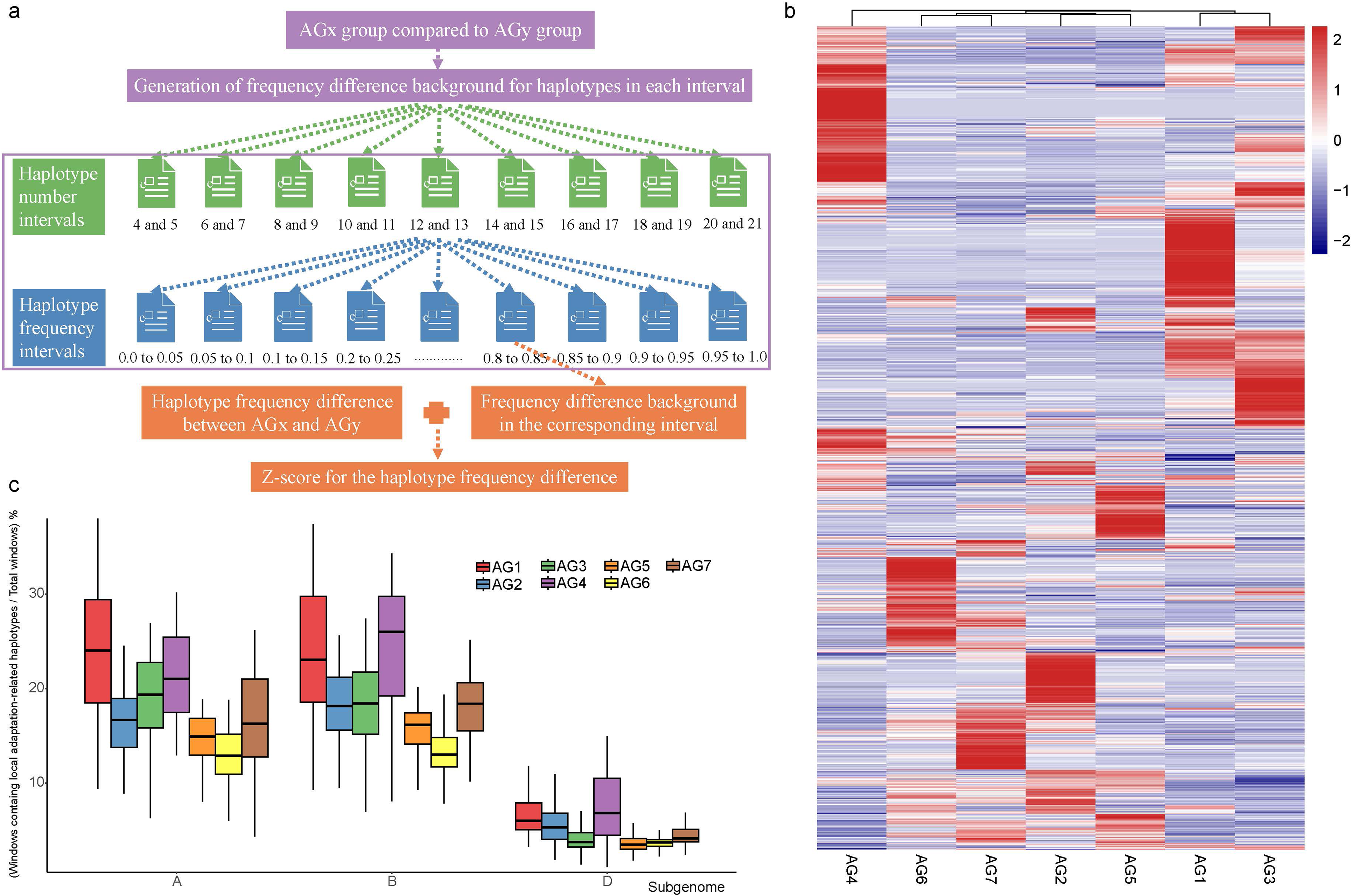
Identification of local adaptation haplotypes (LAr-Haps) for each of the seven Watkins groups. **a**, The diagram showing the haplotype frequency differential-based approach. The genome-wide 1-Mb windows were partitioned into intervals based on the number of haplotypes in each window, and the haplotypes within each window interval were then assigned to 20 intervals according to haplotype frequency. The frequency difference of haplotypes in each interval between the two compared groups was used as the background. The two used intervals reduce the potential adverse effects of large differences in haplotype numbers among windows and in haplotype frequencies. The final Z-score was calculated from the frequency difference of a given haplotype between the two compared groups and the corresponding background. **b**, The frequency comparisons of the identified LAr-Haps among seven Watkins groups. **c**, The percentages of 1-Mb windows containing LAr-Haps for subgenomes A, B, and D. The center line of a boxplot indicates the median; the upper and lower bounds indicate the 75th and 25th percentiles, respectively.

Mapping LAr-Haps onto 1-Mb genomic windows of each of the 827 Watkins accessions, providing a high-resolution map marking the genomic regions related to local environment adaptation^23^, showing that subgenomes A and B harboured substantially more LAr-Haps than subgenome D. The proportion of windows containing LAr-Haps ranged from 12.77% (AG6) to 24.07% (AG1) in subgenome A, 13.32% (AG6) to 24.30% (AG4) in subgenome B, and 3.58% (AG5) to 7.49% (AG4) in subgenome D (Fig. 2c). Distal chromosomal regions were enriched (Supplementary Figs. 7, 8). In several cases, LAr-Haps formed large contiguous blocks, such as a 218-Mb segment on chromosome 6A, spanning from 227 to 445 Mb, detected in 29 AG1 accessions (Supplementary Fig. 9). Notably, these 29 accessions were geographically distinct from the remaining AG1 samples (Supplementary Fig. 10).

Interestingly, we identified a long haplotype block in the corresponding region in our prior study, which was assigned to seven haplotypes (H1-H7) among the 15 pan-genome cultivars representing modern-day diversity across wheat breeding programes^17,30^. This region is associated with several productivity traits (e.g., yield, grain size, height), including the well-known *TaGW2-A* (Grain Width), and was maintained as an intact block in breeding programs to maximize phenotypic expression^30^. To further understand the relationship between the LAr-Haps block in the AG1 group and the seven haplotypes across the 15 wheat cultivars, we first plotted a heatmap of IBSpy values in this region to visualize sequence similarity among related accessions. The results showed 20 of the 29 Watkins accessions and five cultivars (Norin 61, Lancer, Claire, Jagger, SY Mattis) carried the H2 haplotype (Supplementary Fig. 11), suggesting these Watkins accessions as potential donors of this haplotype block. However, the remaining nine accessions carried distinct haplotypes, suggesting the presence of more than one LAr-related long-range haplotype in this region (Supplementary Fig. 12).

### Introgression contributes to the local adaptation of bread wheat

Introgression from wild relatives is a major source of improvement in genetic diversity ^9,10^. To investigate its role in wheat adaptation, we analysed whole-genome resequencing data from 105 wild relatives spanning 43 species (Supplementary Table S4) using the IBSpy pipeline, generating an IBSpy variation matrix^23^. The IBSpy variations clearly separated accessions by ploidy and species in the PCA plot (Supplementary Fig. 13). Applying the IBSpy variation-based introgression detection method developed in our prior study^21^, massive introgressions in the reference genome were determined by detecting the windows with smaller IBSpy variations in wild relatives (IBSpy variations ≤ 30, 40 and 20 for subgenome A, B and D, respectively), including the well-known *A. ventricosa* introgression on chromosome 2A in Jagger (Supplementary Figs. 14 and 15) and *T. timopheevii* introgression on chromosome 2B in LongReach Lancer as positive controls (Supplementary Figs. 16 and 17)^17^. To systematically capture introgressions contained in the Watkins accessions, independent of their presence in the reference genome, we built a 1-Mb haplotype matrix^23^ and applied a minimum tiling path (MTP) strategy developed in our prior study^13^, to identify the longest haplotype matches between Watkins accessions and wild relatives (Supplementary Table S5). In addition to the two introgressions in Jagger and LongReach Lancer that could be accurately detected, some introgressions unique to Watkins accessions or MCs were also identified (Fig. 3a and Supplementary Fig. 18). The length of the introgression fragments contained in each Watkins accession varied among the seven groups, with the average size being 8,658, 8,816, 8,798, 8,951, 8,829, 8,897, 8,885, and 8,718 Mb for AG1 to AG7 and MCs, respectively (Supplementary Fig. 19).

**Fig. 3:**
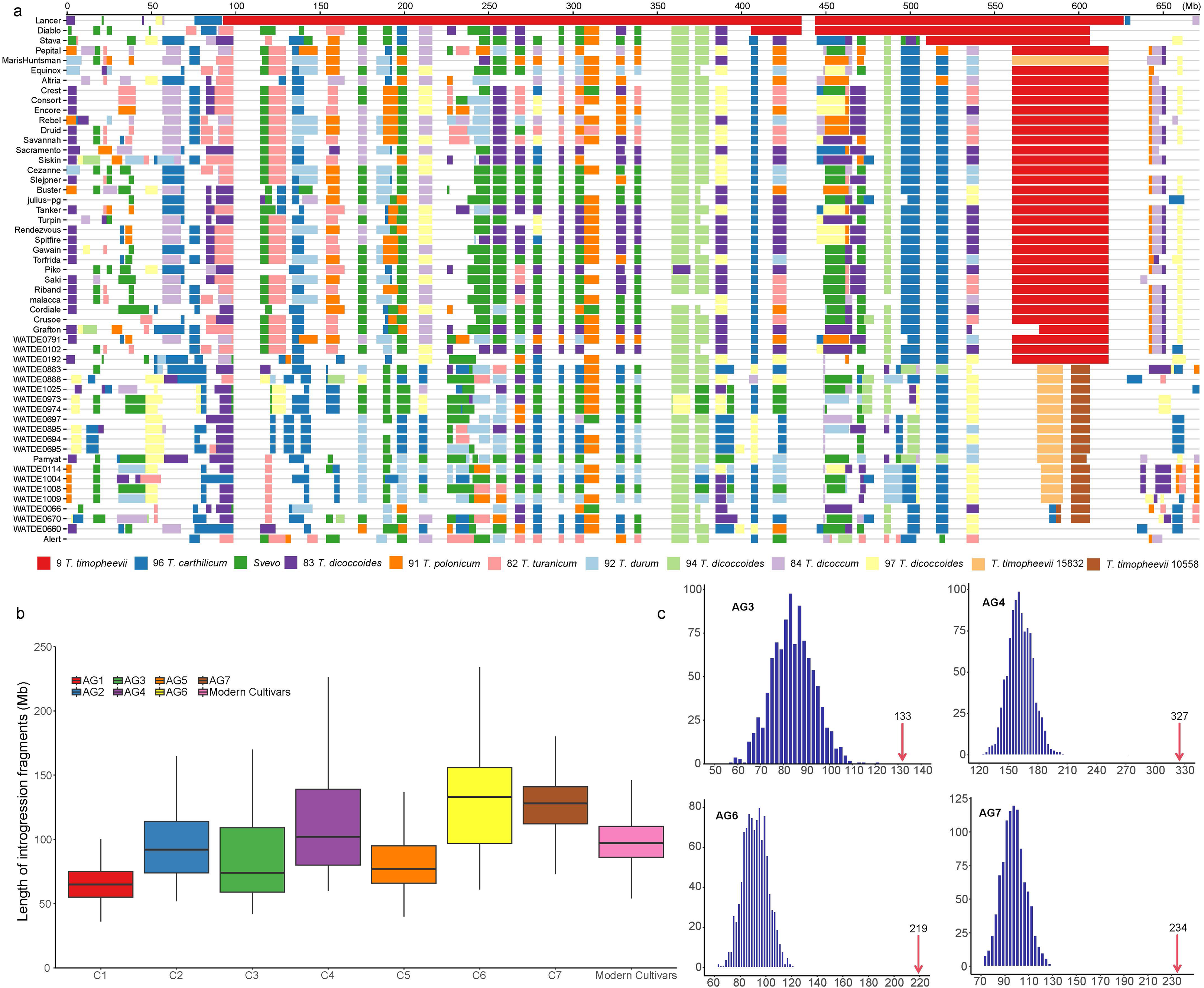
Introgressions from wheat wild relatives and their contributions to local adaptation haplotypes (LAr-Haps). **a**, Diagrammatic representation of *T. timopheevi* introgressions on chromosome 2B in LongReach Lancer, as well as in some Watkins (prefix with “WATDE”) and modern cultivars. The potential donors of the introgressions are listed at the bottom. The detailed information on wild relatives was listed in Supplementary Table 4. The LongReach Lancer genome was used as the reference in this analysis. **b**, Distribution of introgression fragment length in each accession across seven Watkins groups. Only the introgressions from the wild relatives that were further distant from bread wheat (16 AABB genomes were excluded) were counted. The centre line of a boxplot indicates the median; the upper and lower bounds indicate the 75th and 25th percentiles, respectively. **c**, The distribution of LAr-Haps falling within introgressions in the 1,000 permutation tests. For each test, a number of haplotypes equivalent to the number of LAr-Haps in each of the seven Watkins groups was randomly sampled, and the number of sampled haplotypes falling within introgressions was recorded. The observed numbers were marked with red arrows.

Next, we evaluated the contribution of introgressions to the LAr-Haps by aligning the LAr-Haps to the identified introgressions in each Watkins accession. The results revealed that 7,690 (73.16%), 6,220 (74.08%), 6,879 (74.50%), 8,133 (73.13%), 3,887 (75.73%), 5,504 (82.62%), and 6,119 (79.89%) LAr-Haps of AG1 to AG7, respectively, were located in the introgressions of at least one Watkins accession in the corresponding group. We generated a genome-wide background distribution to determine the significance of the observed values through 1,000 permutation tests for each group. In each test, a number of haplotypes equivalent to the number of LAr-Haps was randomly sampled. Given that the number of introgressions varied substantially among the three subgenomes (Supplementary Fig. 20), the randomly sampled haplotypes were required to have the same subgenome origins and frequencies similar to those of LAr-Haps. We then counted the number of these sampled haplotypes falling within introgressions, with average counts for AG1 to AG7 of 8,071, 6,209, 6,914, 8,051, 3,886, 5,273, and 5,956, respectively. These genome-wide background values were comparable to the observed values (Supplementary Fig. 21). However, the number of LAr-Haps located in the introgressions was reduced to 73, 118, 113, 327, 69, 219, and 234 for AG1 to AG7, when restricting analysis to more distantly related wild relatives by excluding the 16 AABB genomes (Fig. 3b and Supplementary Table S6, Fig. 22 and Fig. 23). These genome-wide background values were 103, 111, 85, 163, 59, 92, and 98 for AG1 to AG7, indicating the LAr-Haps of AG3, AG4, AG5, AG6, AG7 were significantly enriched in introgressions, with the enrichment folds being 1.56, 2.01, 1.17, 2.38 and 2.39, respectively (Fig. 3c). These results indicate that introgressions from distant wild relatives have substantially contributed to local adaptation in bread wheat.

### Breeding selection reduces local adaptation haplotype frequencies

In addition to local environmental adaptation, breeding selection has also substantially changed genomes to improve agricultural traits. We identified breeding selection-related haplotypes (BSr-Haps) by comparing haplotype frequencies between MCs and AG2 or AG5, which represent the likely progenitors of European cultivars^13^. We detected 8,177 haplotypes across 5,961 windows (AG2 vs. MC) and 8,854 haplotypes across 5,895 windows (AG5 vs. MC), with significant frequency differences (Supplementary Tables S7, S8). Many genes that have been experimentally verified to function in regulating plant architecture, yield components, growth period, end-use quality, and resistance to foliar diseases were located in the identified windows (Fig. 4a and Supplementary Fig. 24). Most BSr-Haps decreased in frequency in MCs relative to LAs: 6,891 (84.27%) in AG2 comparisons and 5,973 (67.46%) in AG5, both significantly greater than genome-wide expectations from permutation tests, 5,545 (67.81%) for AG2 and 5,467 (61.75%) for AG5 group, respectively. This suggests that modern breeding often dilutes haplotypes prevalent in landraces.

**Fig. 4:**
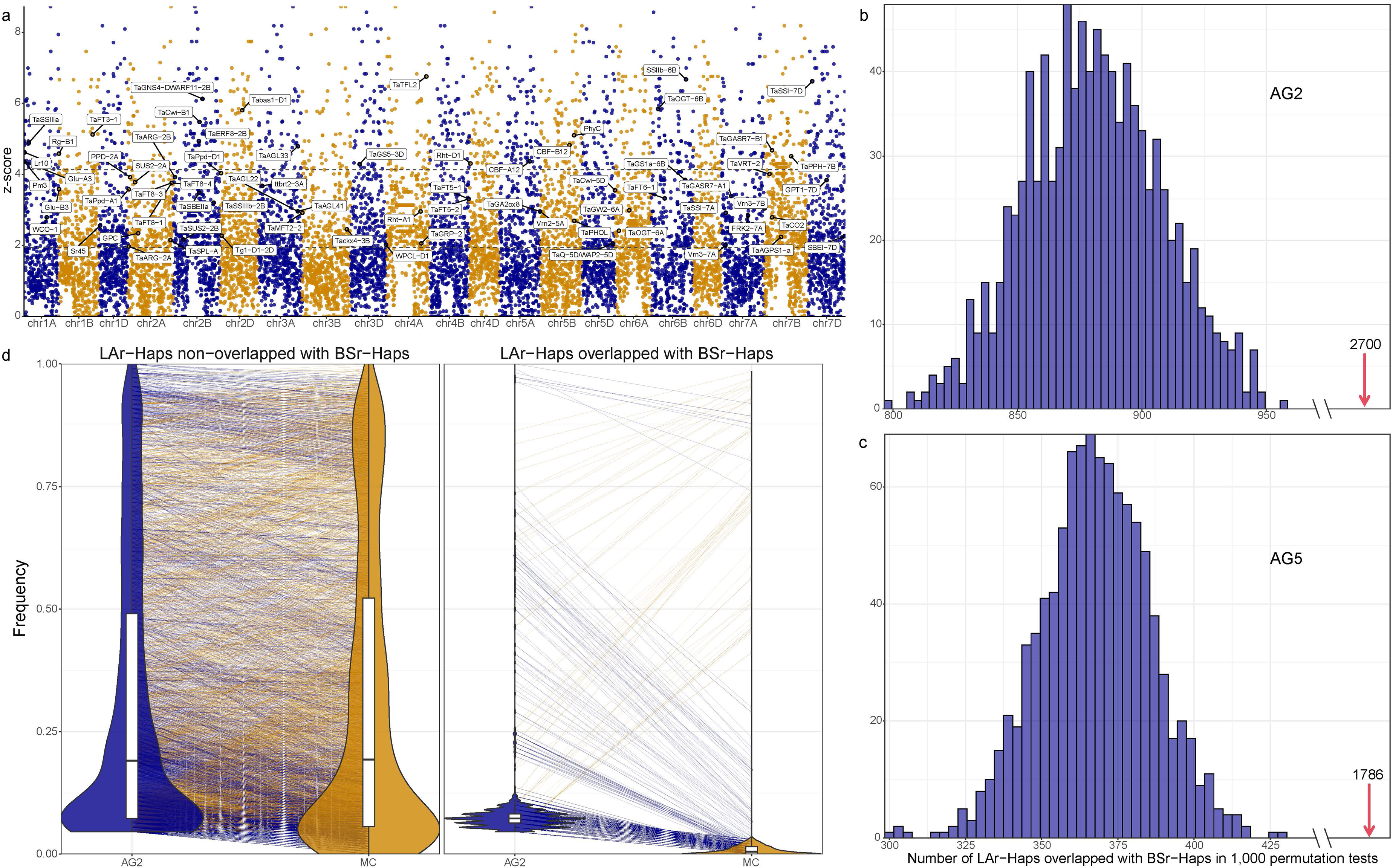
Identification of breeding selection-related haplotypes (BSr-Haps) and frequency change of BSr-Haps between Watkins and modern cultivars. **a**, Distribution of Z-scores of BSr-Hap frequency difference between AG2 and modern cultivars. The two horizontal dashed lines indicate the cut-offs of the top 1% and 5% of Z-scores. Known genes that overlap with windows containing BSr-Haps are marked. **b-c**, Comparison of the observed number of overlapped local adaptation-related haplotypes (LAr-Haps) and BSr-Haps in AG2 (**b**) and AG5 (**c**) to the corresponding genome-wide background numbers generated by 1,000 permutation tests. In each test, a number of haplotypes equivalent to the number of LAr-Haps was randomly sampled, and the number of overlapped haplotypes with BSr-Haps was recorded. The observed numbers were marked with red arrows. **d**, The frequency changes of LAr-Haps overlapped (right) and non-overlapped (left) with BSr-Haps. The orange and blue lines indicate the haplotypes that are more and less frequent in AG2 than in modern cultivars.

Next, we aligned the LAr-Haps and BSr-Haps to exploit the relationships between local adaptation and breeding selection. The results revealed 2,700 and 1,786 overlapped haplotypes for AG2 and AG5, which were much larger than the average genome-wide background values 881 and 368 generated by 1,000 permutation tests, representing enrichment folds of 3.06-fold and 4.85-fold, respectively (Fig. 4b,c). This enrichment suggests that local adaptation and breeding selection preferentially target the same haplotypes, posing an intriguing question of whether they drive these overlapping haplotypes in the same direction. By comparing the haplotype frequency between AG2/AG5 and MC, we found that the frequencies of 2,641 (97.81%) and 1,772 (99.22%) overlapped haplotypes in AG2 and AG5, respectively, were declined in MC (Fig. 4c and Supplementary Fig. 25). These results suggested that breeding selection frequently targeted the same haplotypes as natural adaptation but drove them in the opposite direction.

### Genetic effects of the overlapped haplotypes are opposite to the breeding targets

We hypothesized that the reduction of adaptive haplotypes during breeding reflects their unfavourable effects on agronomic traits. To test this, we evaluated the genetic effects of overlapping LAr-Haps and BSr-Haps (2,700 and 1,786 haplotypes in AG2 and AG5, respectively) using 3,545 marker–trait associations from nested association mapping (NAM) populations derived from 73 Paragon × Watkins crosses, evaluated for 137 traits across 10 environments over 10 years^13^. Taking AG2 as an example, 839 (31.07%) overlapping haplotypes segregated between parents in seven ‘Paragon’ × Watkins recombinant inbred line (RIL) populations. Of these, 312 haplotypes were located within the identified QTL intervals controlling 20 agricultural traits, indicating that 37.20% of the evaluated haplotypes had genetic effects on these traits (Supplementary Tables S9 and S10). To determine the percentage in the genome-wide background, we performed 1,000 permutation tests by randomly sampling 2,700 haplotypes from the AG2 group and recording the number of haplotypes with genetic effects on agricultural traits in each test. This analysis revealed a percentage ranging from 48.70% to 41.9%, with an average of 45.28%, significantly larger than the observed value of 37.20% (Fig. 5a, b). This suggests that overlapping haplotypes are less frequently associated with typical agronomic traits than random haplotypes.

**Fig. 5:**
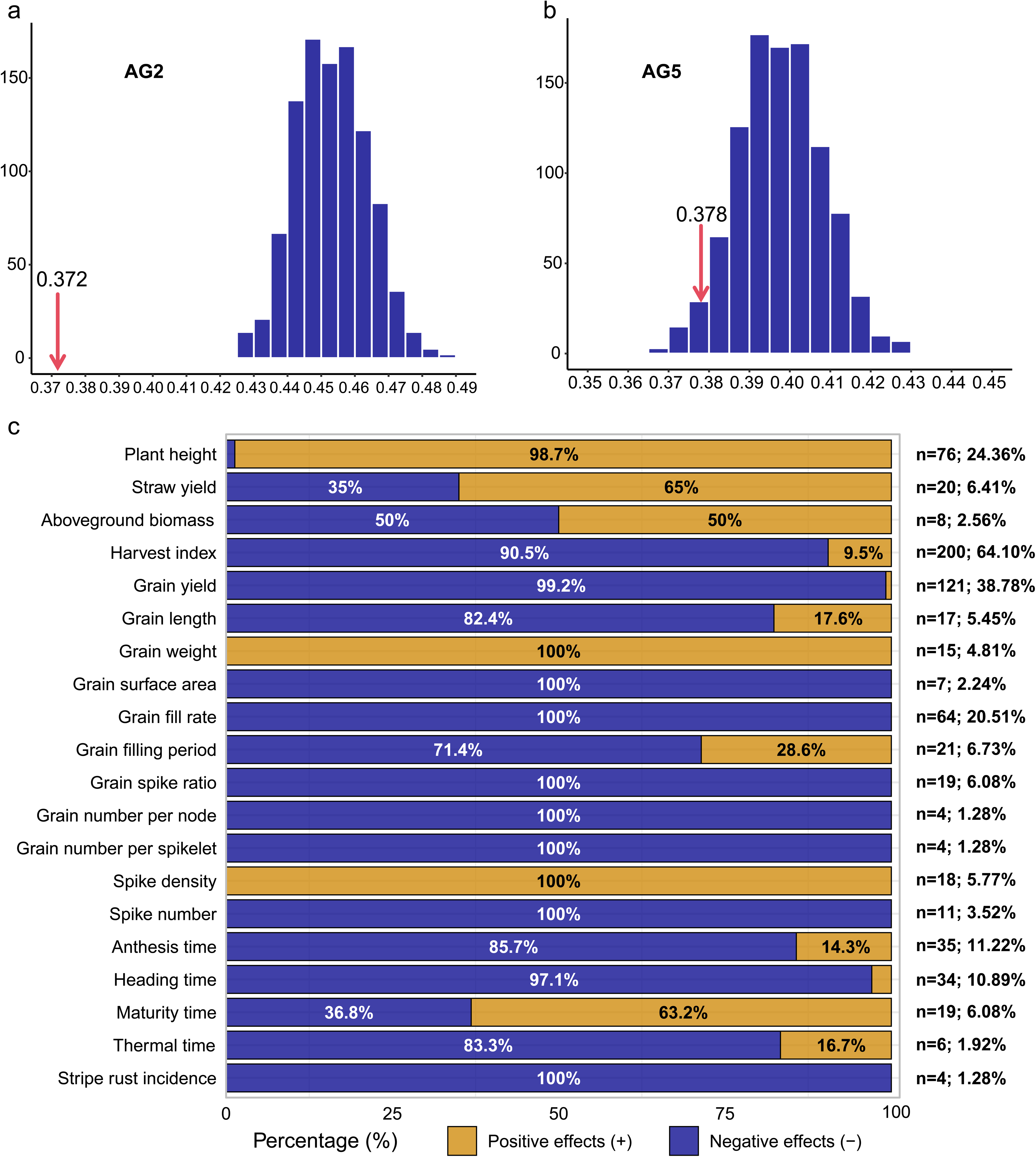
Genetic effects of the overlapped haplotypes between local adaptation-related haplotypes (LAr-Haps) and breeding selection-related haplotypes (BSr-Haps) on agricultural traits. **a**, Comparison of the ratio of overlapped haplotypes having genetic effects on agricultural traits to the genome-wide background values generated from 1,000 permutation tests. In each test, 2,700 (AG2) and 1,786 (AG5) haplotypes were randomly sampled, and the number of haplotypes with genetic effects on agricultural traits was recorded for each test. The observed numbers were marked with red arrows. **b**, The percentages of overlapped haplotypes having positive and negative effects on each of the investigated agricultural traits in AG2. The positive or negative effect indicates that the phenotype values of recombinant inbred lines harbouring overlapped haplotype are higher or lower than those of lines harbouring alternative haplotype in the corresponding recombinant inbred line population. “n” and percentages on the right indicates the numbers of the overlapped haplotypes having genetic effects on the corresponding agricultural traits, and the percentages of these haplotypes to the total overlapped haplotypes having genetic effects on agricultural traits in AG2, respectively.

Next, the genetic effects of the overlapping haplotypes on agricultural traits were categorized as positive or negative by comparing trait values between lines harbouring alternative haplotypes (positive/negative effect: lines harbouring the target haplotypes show higher/lower trait values). Notably, the genetic effect categories of most overlapping haplotypes were unfavourable in wheat breeding (Fig. 5c and Supplementary Fig. 26). For example, 75 (98.70%) of the 76 haplotypes having genetic effects on plant height showed positive effects, meaning these haplotypes increased plant height compared to the alternative haplotypes. Additionally, 99.2%, 100%, and 90.5% of the 121, 64, and 200 haplotypes, respectively, that have genetic effects on grain yield, grain filling rate, and harvest index, showed negative effects. These results explain why adaptive haplotypes were preferentially excluded during breeding: their effects on agronomic traits conflicted with modern breeding targets.

## Discussion

Our study presents a haplotype-centred framework for understanding how the interplay between natural and breeding selection has shaped wheat genomes. By analysing a large resequencing dataset of landraces and modern cultivars with an alignment-free haplotype detection approach, we identified local adaptation haplotypes that have contributed to wheat expansion across diverse agroecological zones. At the same time, we showed that breeding selection often acted on the same haplotypes but drove them in opposite evolutionary directions. These findings underscore that adaptive haplotype contributed to environmental resilience while also becoming central to the trade-offs encountered during modern breeding.

Previous research has extensively documented signatures of local adaptation in wheat and its wild relatives, emphasizing the role of natural selection in shaping allelic diversity along environmental gradients^3,9,13,15,31,32^. In contrast, studies explicitly linking natural and breeding selection have been limited. Our results address this gap by demonstrating that local adaptation haplotypes and breeding selection-related haplotypes overlap extensively, but their evolutionary trajectories diverge depending on their effects on agronomic traits. In particular, haplotypes that enhanced adaptation were frequently associated with reduced yield or unfavourable plant architecture, leading to their exclusion from modern cultivars. This perspective bridges ecological adaptation research and breeding-focused studies, underscoring the importance of integrating both viewpoints.

The patterns we observed can be partly explained by the evolutionary history of adaptive haplotypes. Introgressions from distantly related wild relatives often retain rare alleles conferring resilience to environmental stress, even when associated with negative effects on productivity^10,17,21,33,34^. Similarly, the relatively low proportion of overlapping haplotypes that directly affect agronomic traits suggests that variants conferring ecological fitness are not always aligned with the targets of breeding programs. These dynamics illustrate how long-term adaptation processes may conflict with short-term breeding goals, helping explain why many adaptive haplotypes have been filtered out during crop improvement.

Looking forward, disentangling beneficial adaptive variants from linked deleterious effects will be essential for future breeding. High-throughput phenotyping platforms, combined with genomic datasets, could help reveal how specific adaptive haplotypes contribute to environmental resilience, thereby identifying adaptive alleles with high breeding potential^10,35–37^. Moreover, a deeper mechanistic dissection of how certain LAr-Haps negatively affect agricultural traits will be essential for decoupling adaptation from yield penalties^35,38^. Emerging technologies such as gene editing offer opportunities to break unfavourable linkages, enabling the direct use of adaptive haplotypes in breeding. Together, these strategies provide a roadmap for leveraging haplotype diversity to design wheat cultivars that are both resilient and high-yielding under future climate challenges.

## Methods

### Generation of IBSpy matrix and assignment of haplotypes

The IBSpy matrix and the haplotypes were generated with IBSpy (https://github.com/Uauy-Lab/IBSpy), a *k*-mer-based pipeline that counts variations in 50-Kb windows described in our previous studies^13,21^. Briefly, KMC3^39^ was used to build a *k*-mer (*k*_=_31) database from the re-sequenced reads of each of the 1,051 wheat accessions and each of the ten genome assemblies of wheat^17^. Then, the *k*-mers from each 50-Kb non-overlapping window of a given reference genome were compared to the *k*-mer database of a wheat accession, and the number of variations within each window was counted using the IBSpy pipeline. A variation is defined as a set of continuous, overlapping k-mers that are completely absent from the compared wheat accession. This analysis generated an IBSpy variation matrix for each reference genome, indicating the variations across all used wheat accessions relative to that reference genome at 50-Kb resolution. Low IBSpy variation indicates high similarity between the 50-Kb reference sequence and the compared wheat accession, whereas high IBSpy variation indicates lower sequence similarities.

Next, the IBSpy variations from 20 consecutive 50-Kb windows were extracted for each wheat accession, and the 20 IBSpy variations from all 1,051 wheat accessions were compiled into a matrix (one line per wheat accession), which was then passed to the cluster analysis using the affinity propagation clustering technique^40^. Accessions from the same cluster were assigned to the same haplotype. To enrich the IBSpy variations used for clustering, the IBSpy values derived from syntenic regions across the 10+ Wheat Genomes Project references were incorporated. This analysis generated a haplotype matrix for each reference genome, indicating the haplotypes of all used wheat accessions at 1-Mb resolution (20 consecutive 50-Kb windows).

### Identification of LAr-Haps and BSr-Haps

The LAr-Haps and BSr-Haps were identified based on the haplotype frequency difference in the two compared groups, with the following three steps:

1. The haplotype window filtering. To reduce the potential noise induced by the outlier windows (1-Mb), which harboured too many or too few haplotypes, the number of haplotypes in each window was checked, and the windows in which the number was in the top 5% or the bottom 5% among genome-wide windows were removed from the following analysis.
2. Generation of the genome-wide background of haplotype frequency. The interval set of haplotype numbers was first generated with a step size of 2, and then the chromosome windows were assigned to the corresponding interval based on the number of haplotypes in each window. For each haplotype number interval, the interval set of haplotype frequencies was generated with a step size of 0.05, resulting in 20 intervals. The frequency ranges of the first and last intervals were from 0.00 to 0.05 and from 0.95 to 1.00, respectively. All haplotypes from the windows within each haplotype-number interval were assigned to the frequency interval based on their frequencies in the two compared groups. Then, for each haplotype within the given frequency interval, the frequency difference between the two compared groups was calculated, and the average and standard deviation were computed for each interval, which served as the background for the frequency differences of haplotypes with similar features.
3. Production of z-scores for the frequency difference of each haplotype. Genome-wide haplotypes were scanned individually, and the frequency difference between the two groups was calculated. For a given haplotype, the corresponding haplotype number and haplotype frequency interval were determined, and the average and standard deviation values were extracted. Then, the z-score was calculated using the haplotype frequency difference, the average, and the standard deviation. The haplotypes with the top 5% z-scores were considered differential at the frequency level.
4. Each pair of the seven groups (AG1 to AG7) was compared with the above strategy, and the haplotypes with significantly different frequencies were defined as LAr-Haps. For breeding selection, haplotype frequencies in MCs were compared with those in AG2 and AG5, which represent their ancestral groups^13^, and haplotypes showing significant differences were defined as BSr-Haps.

### Introgression detection

Building upon the assigned haplotypes in non-overlapping 1-Mb windows, introgressions in each Watkins accession were detected using extended matched haplotype blocks from the wheat wild relatives, employing an in silico minimum tiling path approach (https://github.com/Uauy-Lab/MLP_finding) developed in our previous study^13^. Briefly, for each Watkins accession, the haplotypes per chromosome were compared to the corresponding chromosome’s haplotypes of each Watkins line. The length of matched haplotype windows (MHWs) from each wild relative was noted, and only the wild relative with the longest MHW (minimum length was set to three continuous windows) was retained. Then, the comparison was iterated from the second window onward, stepping to the next window until it covered the entire chromosome. In each comparison, the longest MHW and its corresponding wild relative were recorded. Next, the identified MHWs from each window were aligned to the chromosome by their physical positions, and those that were completely covered by others were removed. This approach identified the closest wild relatives at a 1-Mb resolution for any given genomic region within a Watkins accession. The MHWs within the Watkins accession were regarded as introgression fragments contributed by the associated wild relative, because they harboured identical haplotypes in the MHWs.

## Acknowledgements

This work was supported by the Biological Breeding-National Science and Technology Major Project (No. 2023ZD0402403 to Xiaoming Wang and No. 2023ZD0402401 to Shengbao Xu). We thank the support of NWAFU and the John Innes Centre’s High-Performance Computing.

## Author contributions

C.U. and X.W. conceived the project and designed the research; X.W. performed the data analysis; J.QC and R.H. RZ developed the IBSpy, and J.QC developed the affinity propagation; Z.X., X.S. (Xue Shi), and X.S. (Shengbao Xu) analysed the relationship between selected haplotypes and QTLs regulating agricultural traits; S.A. assembled the panel of wild relatives and delivered the DNA; S.C. performed the sequencing of all used varieties; N.A. helped assemble the variety panel. X.W. and C.U. organised the data and wrote the manuscript.

---

**Fig. S1: Identification of landrace- and modern cultivar-private haplotypes using *k*-mer-based IBSpy variations and haplotypes with Jagger as reference. a**, PCA (Principal Component Analysis) plot of the first two components calculated using the genome-wide IBSpy variations of 827 landraces and 244 modern cultivars. The seven groups of Watkins and modern cultivars were defined in our previous study^13^, and the 15 previously described wheat cultivars from the 10+ Wheat Genomes Project^17^ were highlighted. **b**, Distribution of haplotype number per Mb across all 21 chromosomes (positions scaled to % of maximum chromosome length). Boxplots show distributions of 1% bins. The four chromosomal compartments were defined based on the recombinations^18^. **c**, Number of landraces (Watkins)- and modern cultivar-private haplotypes and the shared haplotypes between these two groups across all 21 chromosomes.

**Fig. S2: Distribution of private haplotypes along the 21 chromosomes.** Heatmap representing the number of private haplotypes in each 1-Mb window. The top and bottom lines for each chromosome indicate the Watkins- and modern-cultivar-private haplotypes, respectively.

**Fig. S3: Diagram showing the private haplotypes of each accession within the Watkins AG1 group along the 21 chromosomes.** The 1-Mb windows harbouring Watkins-private haplotypes were marked with purple for each accession. Chinese Spring was used as the reference in this analysis.

**Fig. S4: Diagram showing the private haplotypes of each modern cultivar along the 21 chromosomes.** The 1-Mb windows harbouring modern cultivar-private haplotypes were marked with purple for each accession. Chinese Spring was used as the reference in this analysis.

**Fig. S5: Diagram showing the private haplotypes of each modern cultivar along the 21 chromosomes.** The 1-Mb windows harbouring modern cultivar-private haplotypes were marked with purple for each accession. Jagger was used as the reference in this analysis.

**Fig. S6: The sharing of local adaptation-related haplotypes across seven Watkins groups.** Various combinations, or those in isolation, are shown in the UpSet plot, ordered by frequency.

**Fig. S7: Distribution of local adaptation-related haplotype number per Mb across all 21 chromosomes (positions scaled to % of maximum chromosome length).** Boxplots show distributions of 1% bins. The four chromosomal compartments were defined based on the recombinations^18^.

**Fig. S8: Distribution of local adaptation-related haplotypes along the 21 chromosomes.** Heatmap representing the number of local adaptation-related haplotypes in each 1-Mb window. From top to bottom for each chromosome, the line indicates the number of local adaptation-related haplotypes of AG1 to AG7, respectively.

**Fig. S9: Diagram showing the local adaptation-related haplotypes of each accession within the Watkins AG1 group along chromosome 6A.** The 1-Mb windows harbouring local adaptation-related haplotypes were marked with purple for each accession. Chinese Spring was used as the reference in this analysis.

**Fig. S10: Geographic distribution of the 59 accessions within the Watkins AG1 group.** “G2” indicates the 29 accessions harbouring a specific genomic block that was composed of continuous LAr-Haps windows, and “G1” indicates the remaining 30 accessions.

**Fig. S11: Diagram showing the sequence similarity among the 59 Watkins AG1 accessions and the cultivars from the 10+ Wheat Genomes Project**^17^ **on the specific genomic block of chromosome 6A.** Sequence similarity was represented by IBSpy values. The IBSpy values were divided into four intervals, as in our other analysis. Specifically, IBSpy <30 on subgenome A (orange windows) indicates a sequence similarity >=99.95%.

**Fig. S12: Haplotypes (1-Mb) of the 59 Watkins AG1 accessions and the cultivars from the 10+ Wheat Genomes Project**^17^ **on the specific genomic block of chromosome 6A.** The numbers in the rectangles differentiate the haplotypes in each 1-Mb window (column).

**Fig. S13: PCA (Principal Component Analysis) plot showing the genetic distance of wild relatives and bread wheat accessions.** The PCA was calculated using the genome-wide IBSpy variations of 105 wild relatives and 11 accessions from the 10+ Wheat Genomes Project^17^. The detailed information on wild relatives was listed in Supplementary Table 4.

**Fig. S14: Sequence similarity of 11 accessions from the 10+ Wheat Genomes Project**^17^ **and some wild relatives on chromosome 2A. Jagger was used as the reference in this analysis.** The sequence similarity was represented by the IBSpy values. The IBSpy values were divided into four intervals from our other analysis. Specifically, IBSpy <=30 on subgenome A (orange windows) indicates a sequence similarity >=99.95%. The detailed information on wild relatives was listed in Supplementary Table 4.

**Fig. S15: Sequence similarity of 11 accessions from the 10+ Wheat Genomes Project**^17^ **and some modern cultivars on chromosome 2A. Jagger was used as the reference in this analysis.** The sequence similarity was represented by the IBSpy values. The IBSpy values were divided into four intervals from our other analysis. Specifically, IBSpy <=30 on subgenome A (orange windows) indicates a sequence similarity >=99.95%. Accessions with orange windows at the start of this chromosome harbour the well-known *A. ventricosa* introgression on chromosome 2A in Jagger, due to sequence similarity with Jagger.

**Fig. S16: Sequence similarity of 11 accessions from the 10+ Wheat Genomes Project**^17^ **and some wild relatives on chromosome 2B. LongReach Lancer was used as the reference in this analysis.** The sequence similarity was represented by the IBSpy values. The IBSpy values were divided into four intervals from our other analysis. Specifically, IBSpy <=40 on subgenome B (orange windows) indicates a sequence similarity >=99.95%. The detailed information on wild relatives was listed in Supplementary Table 4.

**Fig. S17: Sequence similarity of 11 accessions from the 10+ Wheat Genomes Project**^17^ **and some modern cultivars on chromosome 2B. LongReach Lancer was used as the reference in this analysis.** The sequence similarity was represented by the IBSpy values. The IBSpy values were divided into four intervals from our other analysis. Specifically, IBSpy <=40 on subgenome B (orange windows) indicates a sequence similarity >=99.95%. Accessions with orange windows in the middle of this chromosome harbour the well-known *T. timopheevii* introgression on chromosome 2B in LongReach Lancer, due to sequence similarity with LongReach Lancer.

**Fig. S18: Diagrammatic representation of *A. ventricosa* introgression on chromosome 2A in Jagger, as well as in some modern cultivars.** The potential donors of the introgressions are listed at the bottom. The Jagger genome was used as the reference in this analysis.

**Fig. S19: Distribution of introgression fragment length in each accession across seven Watkins groups and modern cultivar.** The centre line of a boxplot indicates the median; the upper and lower bounds indicate the 75th and 25th percentiles, respectively.

**Fig. S20: Distribution of introgression fragment length in each subgenome across seven Watkins groups and modern cultivar.** The centre line of a boxplot indicates the median; the upper and lower bounds indicate the 75th and 25th percentiles, respectively.

**Fig. S21: The distribution of LAr-Haps falling within introgressions in the 1,000 permutation tests.** For each test, a number of haplotypes equivalent to the number of LAr-Haps in each of the seven Watkins groups was randomly sampled, and the number of sampled haplotypes falling within introgressions was recorded, respectively. Given that the number of introgressions varied substantially among the three subgenomes, the randomly sampled haplotypes were required to possess the same subgenome origins and similar frequencies to LAr-Haps. The observed numbers were marked with red arrows.

**Fig. S22: Diagrammatic representation of *T. timopheevi* introgression on chromosome 2B in LongReach Lancer, as well as in some Watkins (prefix with “WATDE”) and modern cultivars.** The potential donors of the introgressions are listed at the bottom. The detailed information on wild relatives was listed in Supplementary Table 4. The LongReach Lancer genome was used as the reference in this analysis. Only the wild relatives that were further distant from bread wheat (16 AABB genomes were excluded) were included in this analysis.

**Fig. S23: Diagrammatic representation of *A. ventricosa* introgression on chromosome 2A in Jagger, as well as in some modern cultivars.** The potential donors of the introgressions are listed at the bottom. The detailed information on wild relatives was listed in Supplementary Table 4. The Jagger genome was used as the reference in this analysis. Only the wild relatives that were further distant from bread wheat (16 AABB genomes were excluded) were included in this analysis.

**Fig. S24: Distribution of Z-scores of frequency difference of breeding selection-related haplotypes (BSr-Haps) between AG5 and modern cultivars.** The two horizontal dashed lines indicate the cut-offs of the top 1% and 5% of Z-scores. Known genes that overlap with windows containing BSr-Haps are marked.

**Fig. S25: The frequency changes of local adaptation-related haplotypes (LAr-Haps) overlapped (right) and non-overlapped (left) with breeding selection-related haplotypes (BSr-Haps).** The orange and blue lines indicate the haplotypes that are more and less frequent in AG5 than in modern cultivars.

**Fig. S26: The percentages of overlapped haplotypes having positive and negative effects on each of the investigated agricultural traits in AG5.** The positive or negative effect indicates that the phenotype values of recombinant inbred lines harbouring overlapped haplotype are higher or lower than those of lines harbouring alternative haplotype in the corresponding recombinant inbred line population. “n” and percentages on the right indicates the numbers of the overlapped haplotypes having genetic effects on the corresponding agricultural traits, and the percentages of these haplotypes to the total overlapped haplotypes having genetic effects on agricultural traits in AG5, respectively.

**Supplementary Table 1: Landrace-private haplotypes.** The column from AG1 to AG7 indicates the number of accessions harbouring the corresponding haplotype in each of the seven Watkins groups. The column of “Modern cultivars” shows the number of accessions harbouring the corresponding haplotype in modern cultivars.

**Supplementary Table 2: Modern cultivar-private haplotypes.** The column from AG1 to AG7 indicates the number of accessions harbouring the corresponding haplotype in each of the seven Watkins groups. The column of “Modern cultivars” shows the number of accessions harbouring the corresponding haplotype in modern cultivars.

**Supplementary Table 3: Local adaptation-related haplotypes (LAr-Haps) in each of the seven Watkins groups.** “T” and “F” indicate whether the haplotype is and is not LAr-Hap in the corresponding Watkins group, respectively.

**Supplementary Table 4: detailed information on wild relatives used in this study.**

**Supplementary Table 5: Introgression windows from wild relatives to Watkins and modern cultivars.** The haplotypes of each query accession were searched against those of wild relatives (as the database) to detect the longest matched haplotype blocks using the MTP (minimum tiling path) strategy. The start and end positions of matched haplotype windows (MHWs) from each query accession were noted.

**Supplementary Table 6: Introgression windows from wild relatives to Watkins and modern cultivars.** The haplotypes of each query accession were searched against those of wild relatives (as the database) to detect the longest matched haplotype blocks using the MTP (minimum tiling path) strategy. The start and end positions of matched haplotype windows (MHWs) from each query accession were noted. Only the wild relatives that were further distant from bread wheat (16 AABB genomes were excluded) were included in this analysis. Detailed information on wild relatives is listed in Supplementary Table 4.

**Supplementary Table 7: Breeding selection-related haplotypes in AG2.** The frequency difference for each haplotype between AG2 and modern cultivars was calculated and compared with the average value and standard deviation in the corresponding background (Fig. 2a), yielding a Z-score to assess significance. The top 5% Z-score threshold is 1.944.

**Supplementary Table 8: Breeding selection-related haplotypes in AG5.** The frequency difference of each haplotype between AG5 and modern cultivars was calculated and compared to the average value and standard deviation in the corresponding background (Fig. 2a), generating a Z-score that indicates significance. The top 5% Z-score threshold is 1.8903.

**Supplementary Table 9: Genetic effects of the overlapped haplotypes in the AG2 analysis on agricultural traits.** The genetic effects were evaluated with 73 ‘Paragon’ × Watkins recombinant inbred lines (RILs). Multiple elements in a single cell are separated by commas.

**Supplementary Table 10: Genetic effects of the overlapped haplotypes in the AG5 analysis on agricultural traits.** The genetic effects were evaluated with 73 ‘Paragon’ × Watkins recombinant inbred lines (RILs). Multiple elements in a single cell are separated by commas.

## References

1 Long, T. et al. The early history of wheat in China from (14)C dating and Bayesian chronological modelling. Nature plants 4, 272–279 (2018). 10.1038/s41477-018-0141-x

2 Smith, O., et al. Archaeology. Sedimentary DNA from a submerged site reveals wheat in the British Isles 8000 years ago. Science (New York, N.Y.) 347, 998–1001 (2015). 10.1126/science.1261278

3 Zhao, X. et al. Population genomics unravels the Holocene history of bread wheat and its relatives. Nature plants 9, 403–419 (2023). 10.1038/s41477-023-01367-3

4 Huang, X. et al. A map of rice genome variation reveals the origin of cultivated rice. Nature 490, 497–501 (2012). 10.1038/nature11532

5 Zhao, Y. et al. The evening complex promotes maize flowering and adaptation to temperate regions. The Plant cell 35, 369–389 (2023). 10.1093/plcell/koac296

6 Lopes, M. S. et al. Exploiting genetic diversity from landraces in wheat breeding for adaptation to climate change. Journal of experimental botany 66, 3477–3486 (2015). 10.1093/jxb/erv122

7 Exposito-Alonso, M., Burbano, H. A., Bossdorf, O., Nielsen, R. & Weigel, D. Natural selection on the Arabidopsis thaliana genome in present and future climates. Nature 573, 126–129 (2019). 10.1038/s41586-019-1520-9

8 Wang, L. et al. The interplay of demography and selection during maize domestication and expansion. Genome Biol 18, 215 (2017). 10.1186/s13059-017-1346-4

9 He, F. et al. Exome sequencing highlights the role of wild-relative introgression in shaping the adaptive landscape of the wheat genome. Nat Genet 51, 896–904 (2019). 10.1038/s41588-019-0382-2

10 Farooq, M. et al. Back into the Wild: Harnessing the Power of Wheat Wild Relatives for Future Crop and Food Security. Journal of experimental botany (2025). 10.1093/jxb/eraf141

11 Allaby, R. G., Ware, R. L. & Kistler, L. A re-evaluation of the domestication bottleneck from archaeogenomic evidence. Evol Appl 12, 29–37 (2019). 10.1111/eva.12680

12 Guo, D. et al. A pangenome reference of wild and cultivated rice. Nature 642, 662–671 (2025). 10.1038/s41586-025-08883-6

13 Cheng, S. et al. Harnessing landrace diversity empowers wheat breeding. Nature 632, 823–831 (2024).

14 Wang, X. et al. HebQTLs reveal intra-subgenome regulation inducing unbalanced expression and function among bread wheat homoeologs. Genome Biol 26, 218 (2025). 10.1186/s13059-025-03694-4

15 Cavanagh, C. R. et al. Genome-wide comparative diversity uncovers multiple targets of selection for improvement in hexaploid wheat landraces and cultivars. Proceedings of the National Academy of Sciences of the United States of America 110, 8057–8062 (2013). 10.1073/pnas.1217133110

16 Hao, C. et al. Resequencing of 145 landmark cultivars reveals asymmetric sub-genome selection and strong founder genotype effects on wheat breeding in China. Molecular plant 13, 1733–1751 (2020).

17 Walkowiak, S. et al. Multiple wheat genomes reveal global variation in modern breeding. Nature 588, 277–283 (2020).

18 IWGSC. Shifting the limits in wheat research and breeding using a fully annotated reference genome. Science (New York, N.Y.) 361 (2018). 10.1126/science.aar7191

19 Jiao, C. et al. Pan-genome bridges wheat structural variations with habitat and breeding. Nature (2024). 10.1038/s41586-024-08277-0

20 Wang, Z. et al. Deciphering the evolution and complexity of wheat germplasm from a genomic perspective. J Genet Genomics 50, 846–860 (2023). 10.1016/j.jgg.2023.08.002

21 Ahmed, H. I. et al. Einkorn genomics sheds light on history of the oldest domesticated wheat. Nature 620, 830–838 (2023). 10.1038/s41586-023-06389-7

22 O’Hara, T. et al. The wheat powdery mildew resistance gene Pm4 also confers resistance to wheat blast. Nature plants 10, 984–993 (2024). 10.1038/s41477-024-01718-8

23. Xiaoming Wang, J. Q.-C., Ricardo H Ramírez-González, Cristobal Uauy. Natural and breeding selection converge on overlapping haplotypes with divergent directions and outcomes in wheat [Data set]. Zenodo 10.5281/zenodo.19257841 (2026).

24 Cheng, H. et al. Frequent intra- and inter-species introgression shapes the landscape of genetic variation in bread wheat. Genome Biol 20, 136 (2019). 10.1186/s13059-019-1744-x

25 Yang, Z. et al. Utilization of 1BL/1RS translocation in wheat breeding in China. Zuo wu xue bao 30, 531–535 (2004).

26 Yang, Z. et al. ggComp enables dissection of germplasm resources and construction of a multiscale germplasm network in wheat. Plant physiology 188, 1950–1965 (2022). 10.1093/plphys/kiac029

27 Gabay, G. et al. Dosage differences in 12-OXOPHYTODIENOATE REDUCTASE genes modulate wheat root growth. Nat Commun 14, 539 (2023). 10.1038/s41467-023-36248-y

28 Charmet, G. Wheat domestication: Lessons for the future. Comptes Rendus. Biologies 334, 212–220 (2011). 10.1016/j.crvi.2010.12.013

29 Igrejas, G. & Branlard, G. in Wheat Quality For Improving Processing And Human Health (eds Gilberto Igrejas, Tatsuya M. Ikeda, & Carlos Guzmán) 1–7 (Springer International Publishing, 2020).

30 Brinton, J. et al. A haplotype-led approach to increase the precision of wheat breeding. Commun Biol 3, 712 (2020). 10.1038/s42003-020-01413-2

31 Zhou, Y. et al. Triticum population sequencing provides insights into wheat adaptation. Nat Genet 52, 1412–1422 (2020). 10.1038/s41588-020-00722-w

32 Wang, H. et al. Sympatric speciation of wild emmer wheat driven by ecology and chromosomal rearrangements. Proceedings of the National Academy of Sciences of the United States of America 117, 5955–5963 (2020). 10.1073/pnas.1920415117

33 Zhou, Y. et al. Introgressing the Aegilops tauschii genome into wheat as a basis for cereal improvement. Nature plants 7, 774–786 (2021). 10.1038/s41477-021-00934-w

34 Cavalet-Giorsa, E. et al. Origin and evolution of the bread wheat D genome. Nature 633, 848–855 (2024). 10.1038/s41586-024-07808-z

35 Mao, H. et al. Wheat adaptation to environmental stresses under climate change: Molecular basis and genetic improvement. Mol Plant 16, 1564–1589 (2023). 10.1016/j.molp.2023.09.001

36 Zhang, Z. et al. An automated root phenotype platform enables nondestructive high-throughput root system architecture dissection in wheat. Plant physiology 198 (2025). 10.1093/plphys/kiaf154

37 Zhang, Z. et al. Integrating high-throughput phenotyping and genome-wide association studies for enhanced drought resistance and yield prediction in wheat. The New phytologist 243, 1758–1775 (2024). 10.1111/nph.19942

38 Xiao, J. et al. Wheat genomic study for genetic improvement of traits in China. Sci China Life Sci 65, 1718–1775 (2022). 10.1007/s11427-022-2178-7

39 Kokot, M., Dlugosz, M. & Deorowicz, S. KMC 3: counting and manipulating k-mer statistics. Bioinformatics 33, 2759–2761 (2017). 10.1093/bioinformatics/btx304

40 Frey, B. J. & Dueck, D. Clustering by passing messages between data points. Science (New York, N.Y.) 315, 972–976 (2007). 10.1126/science.1136800

