## Supplementary figures and images for "Natural and breeding selection converge on overlapping haplotypes with divergent directions and outcomes in wheat"

### Fig. S1.pdf

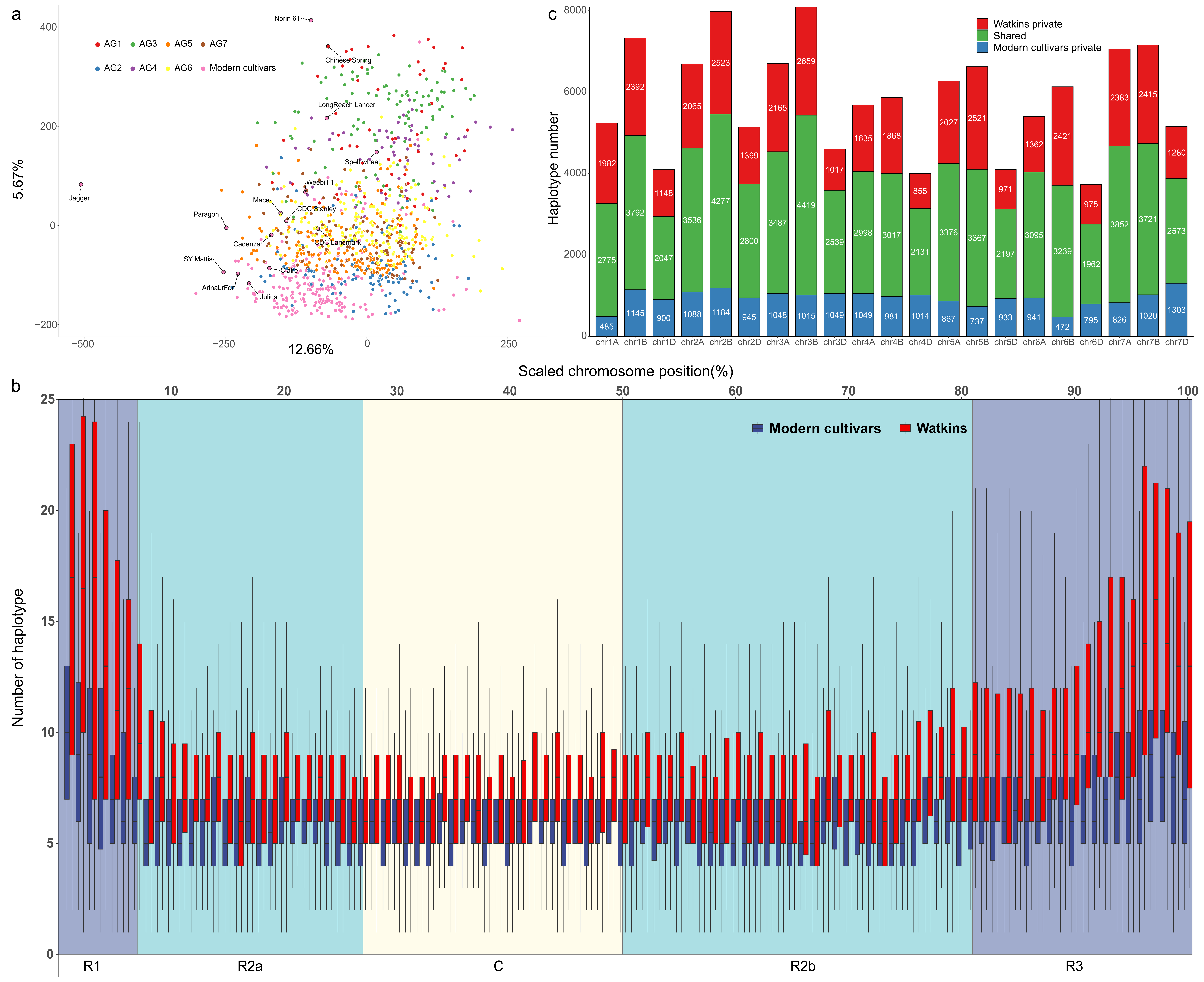

### Fig. S2.pdf

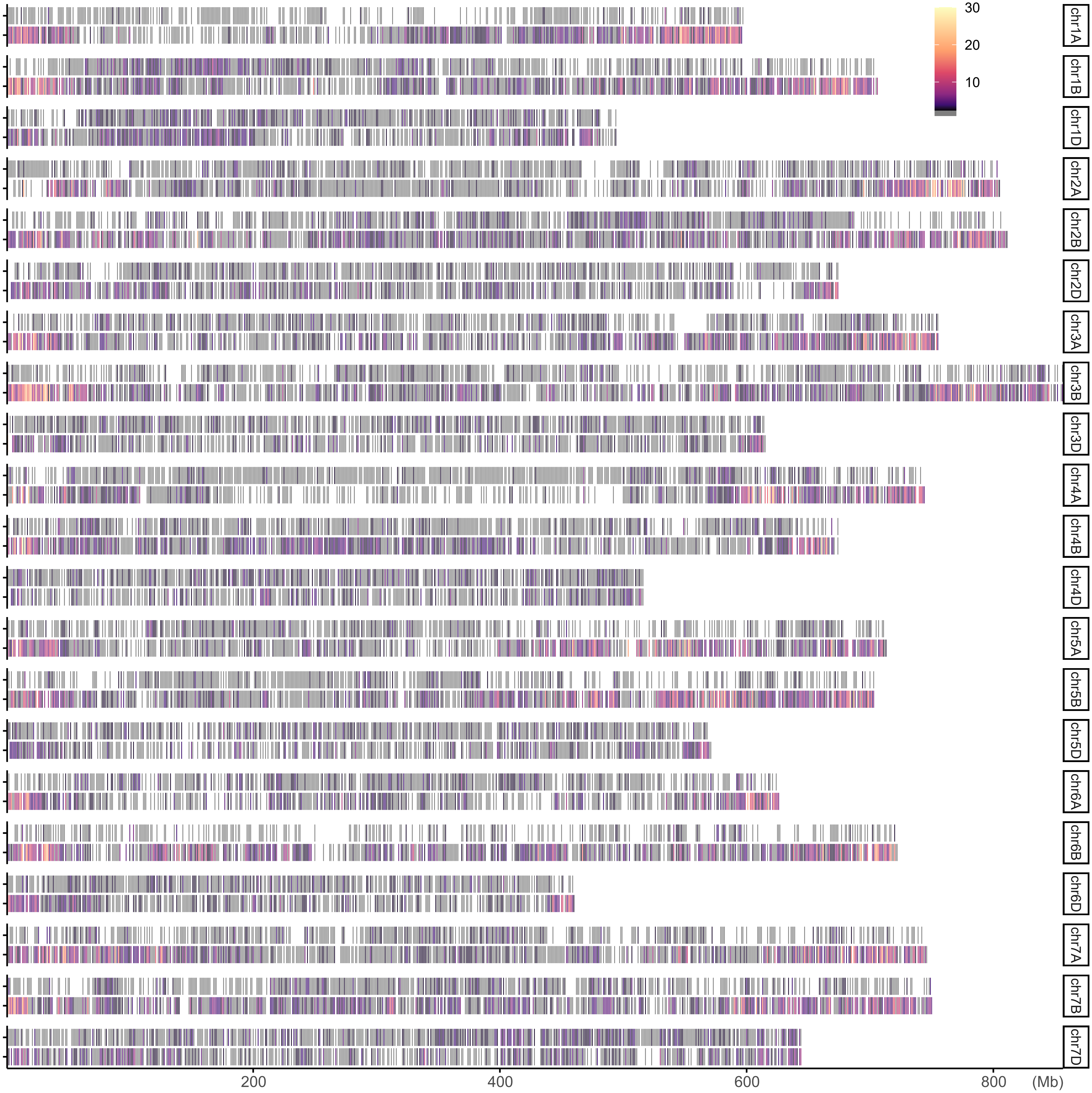

### Fig. S3.pdf

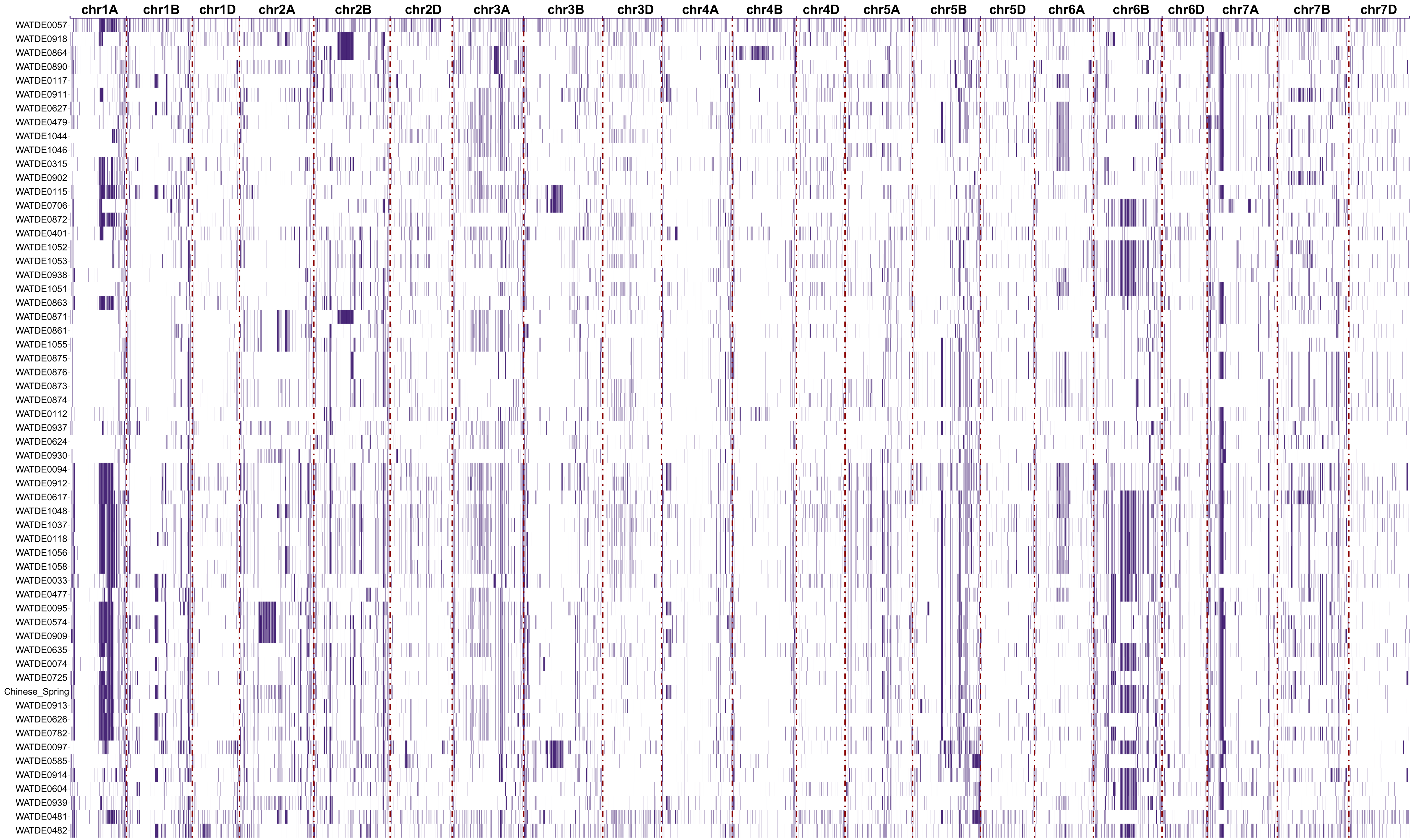

### Fig. S4.pdf

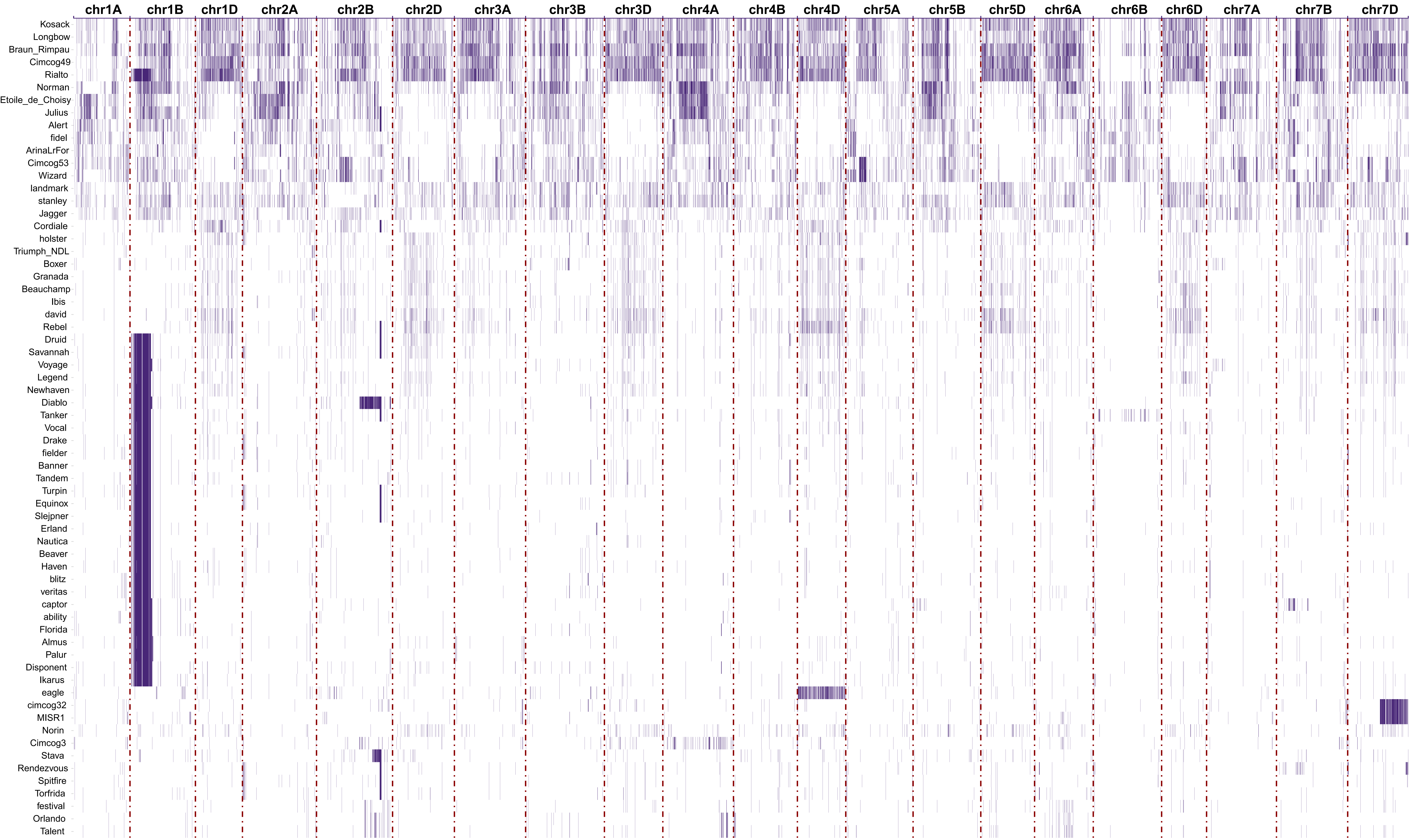

### Fig. S5.pdf

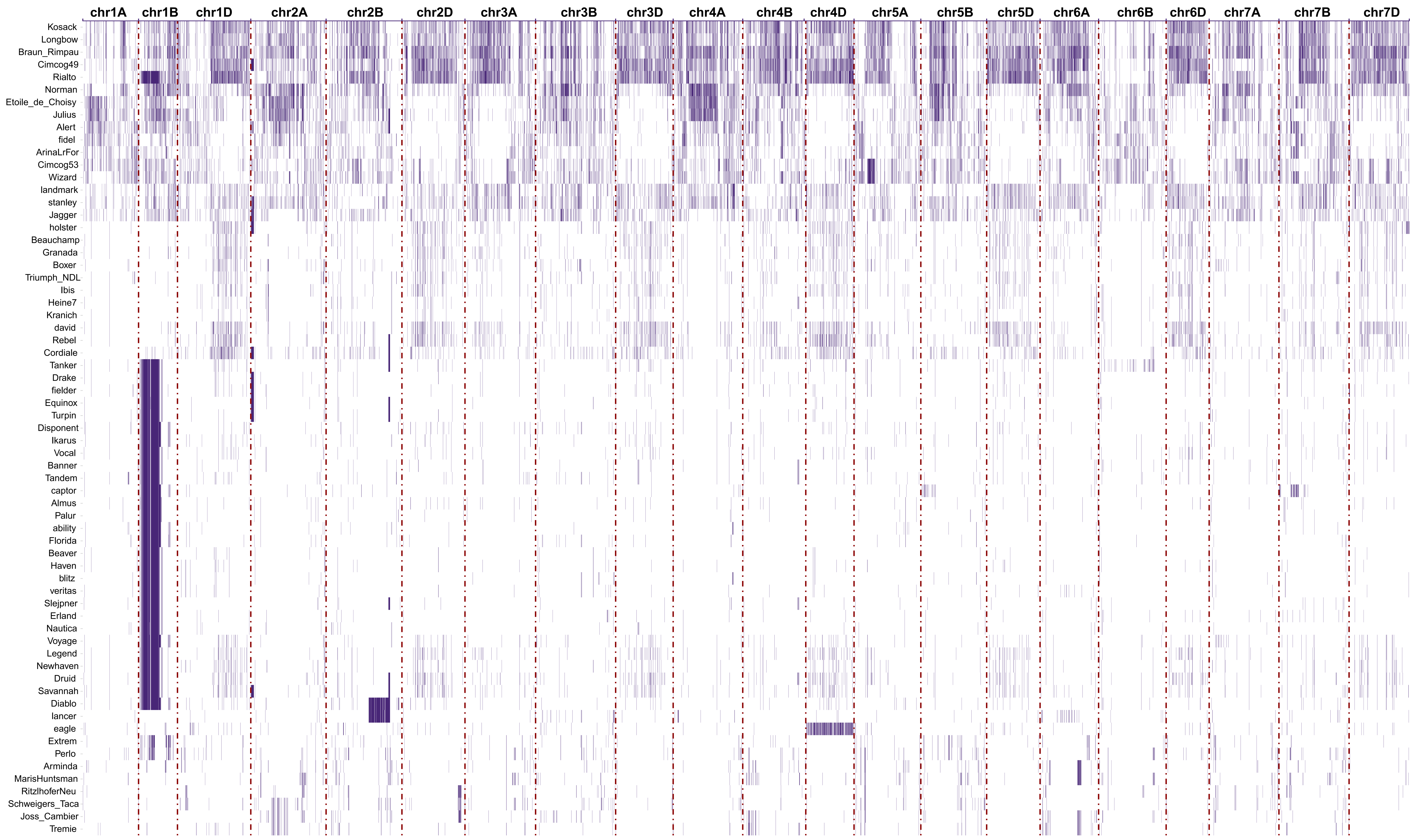

### Fig. S6.pdf

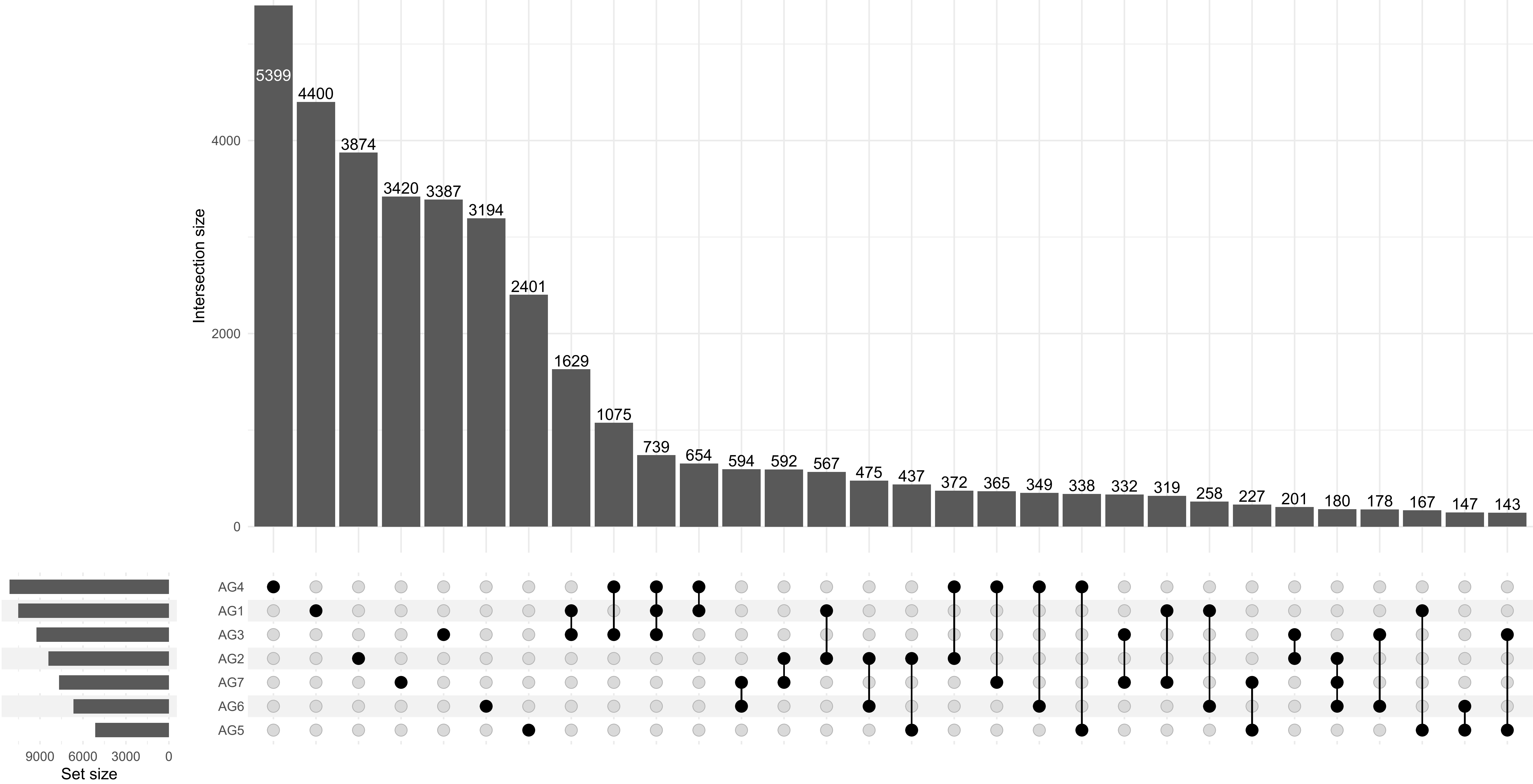

### Fig. S7.pdf

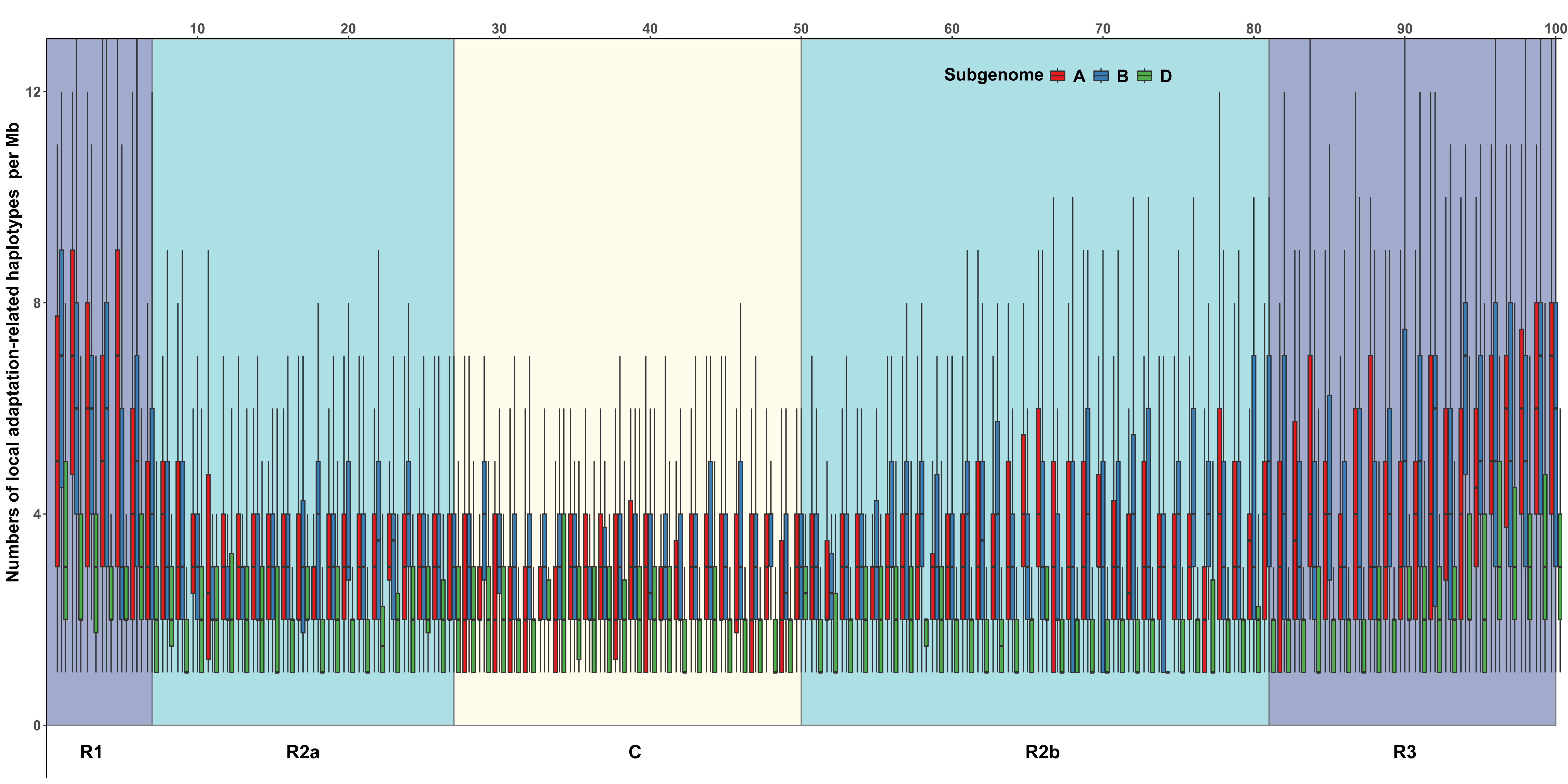

### Fig. S8.pdf

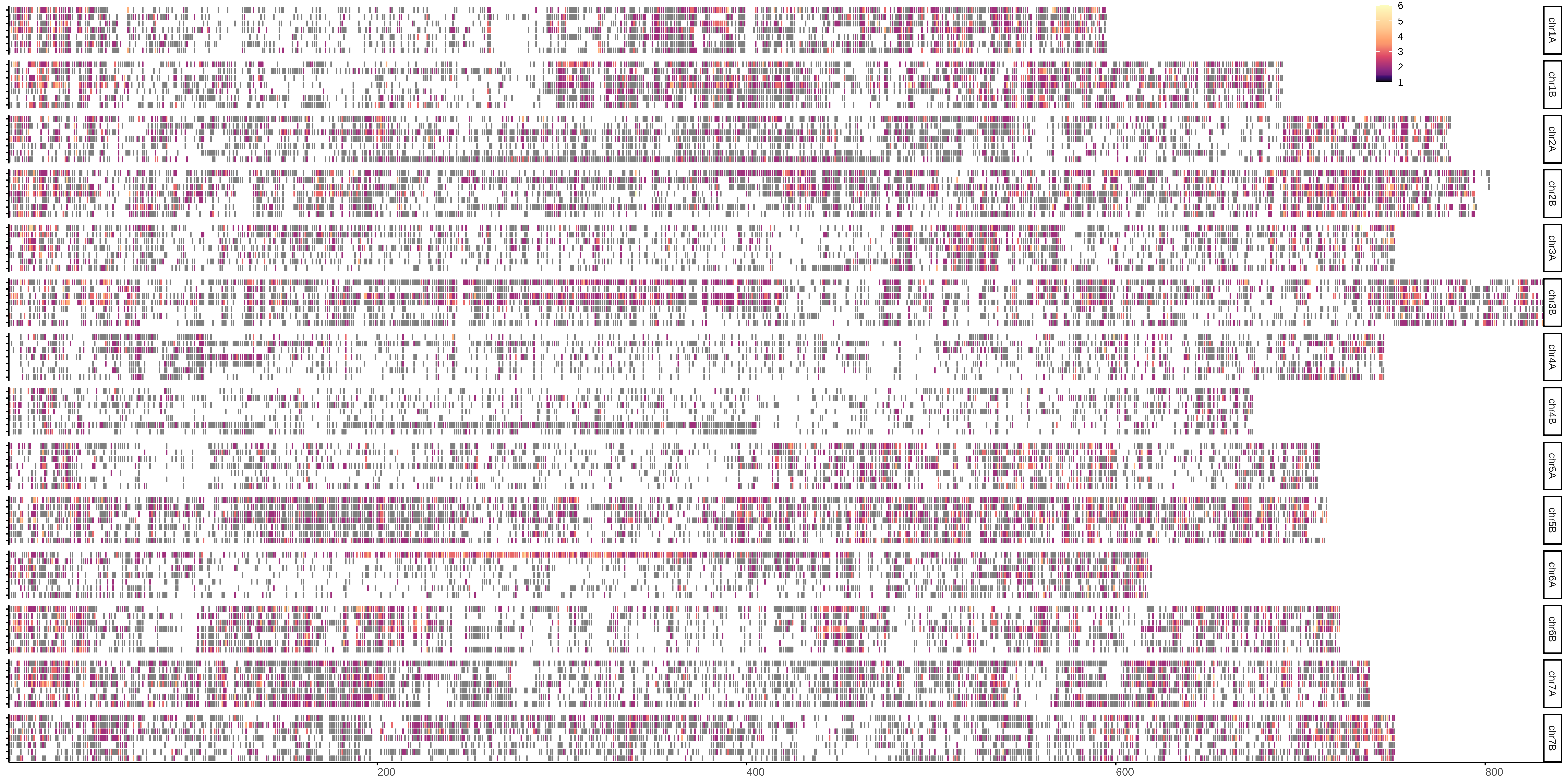

### Fig. S9.pdf

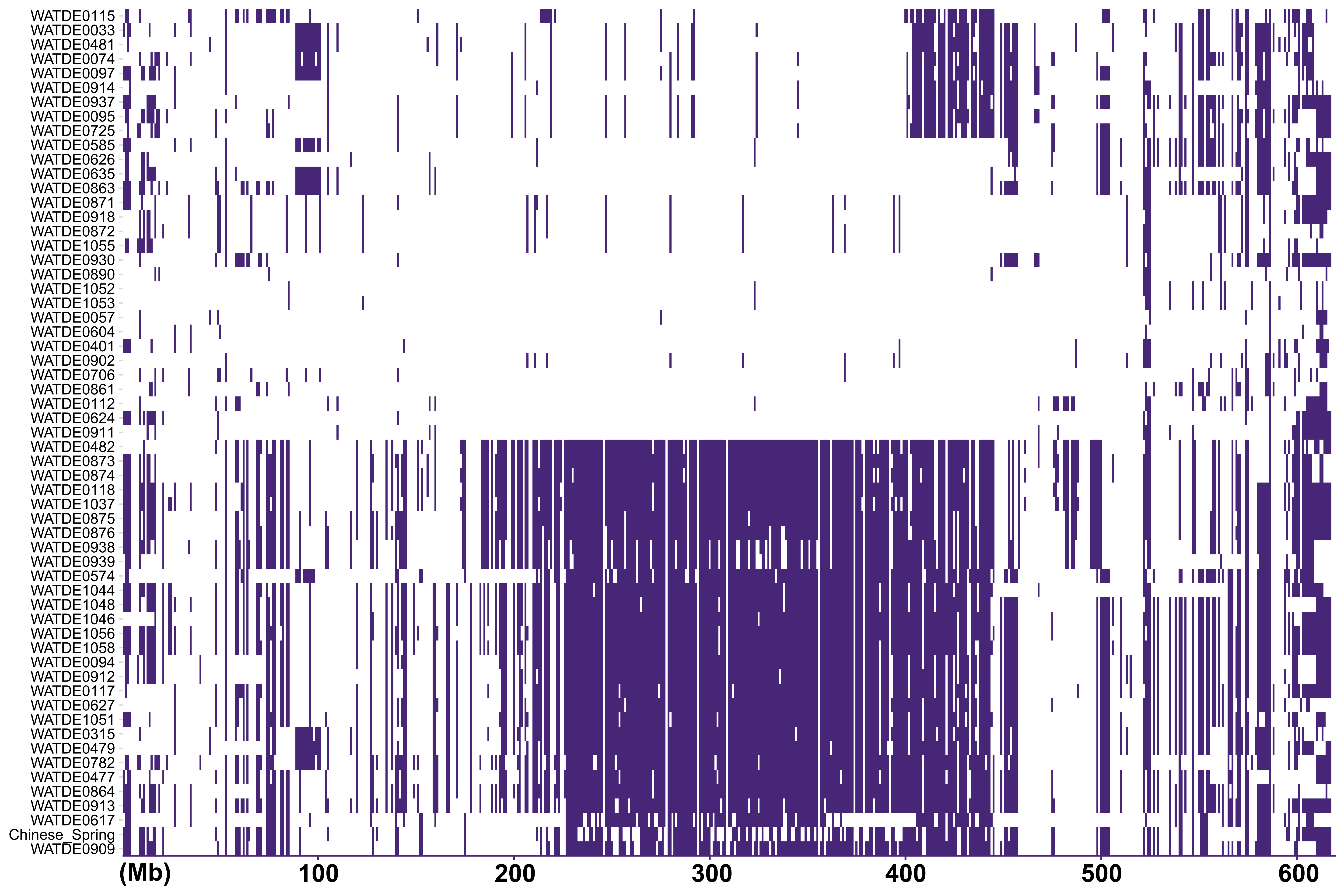

### Fig. S10.pdf

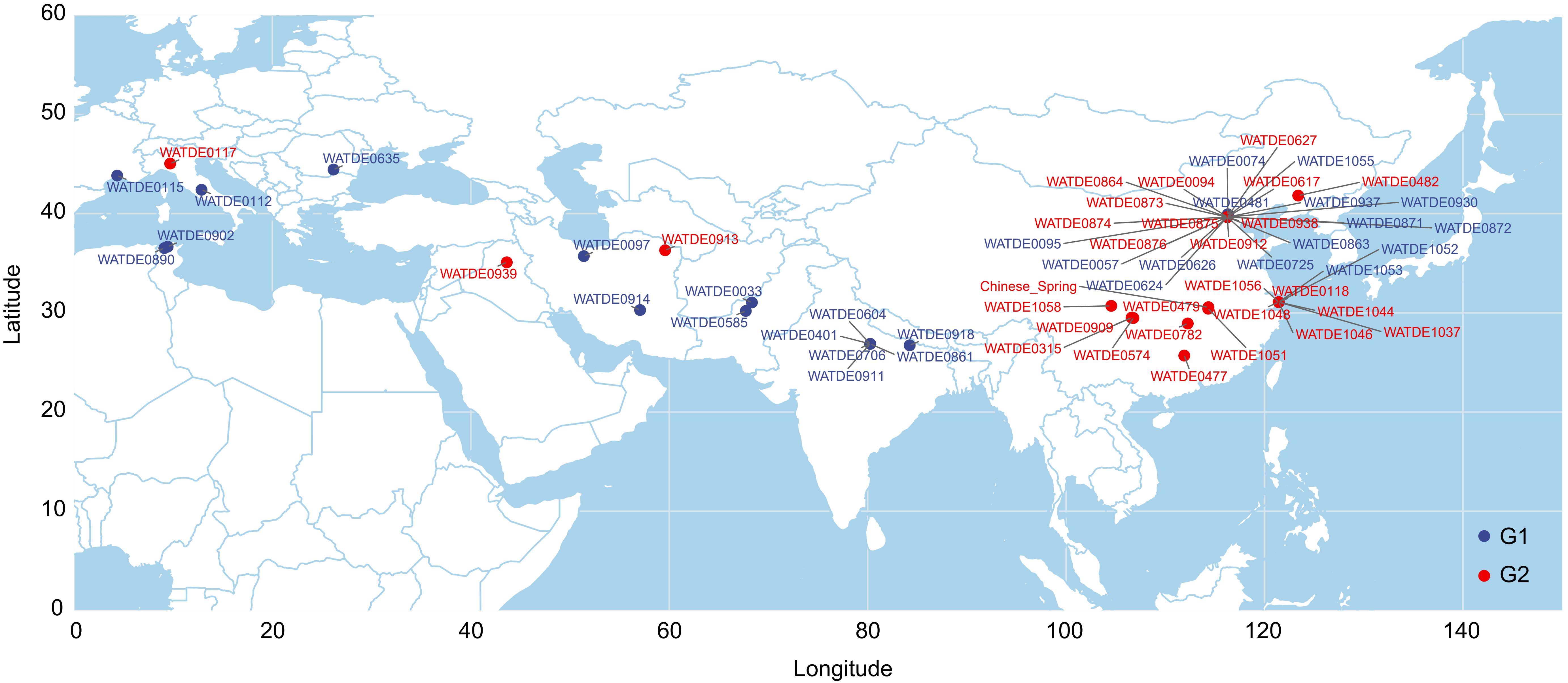

### Fig. S11.pdf

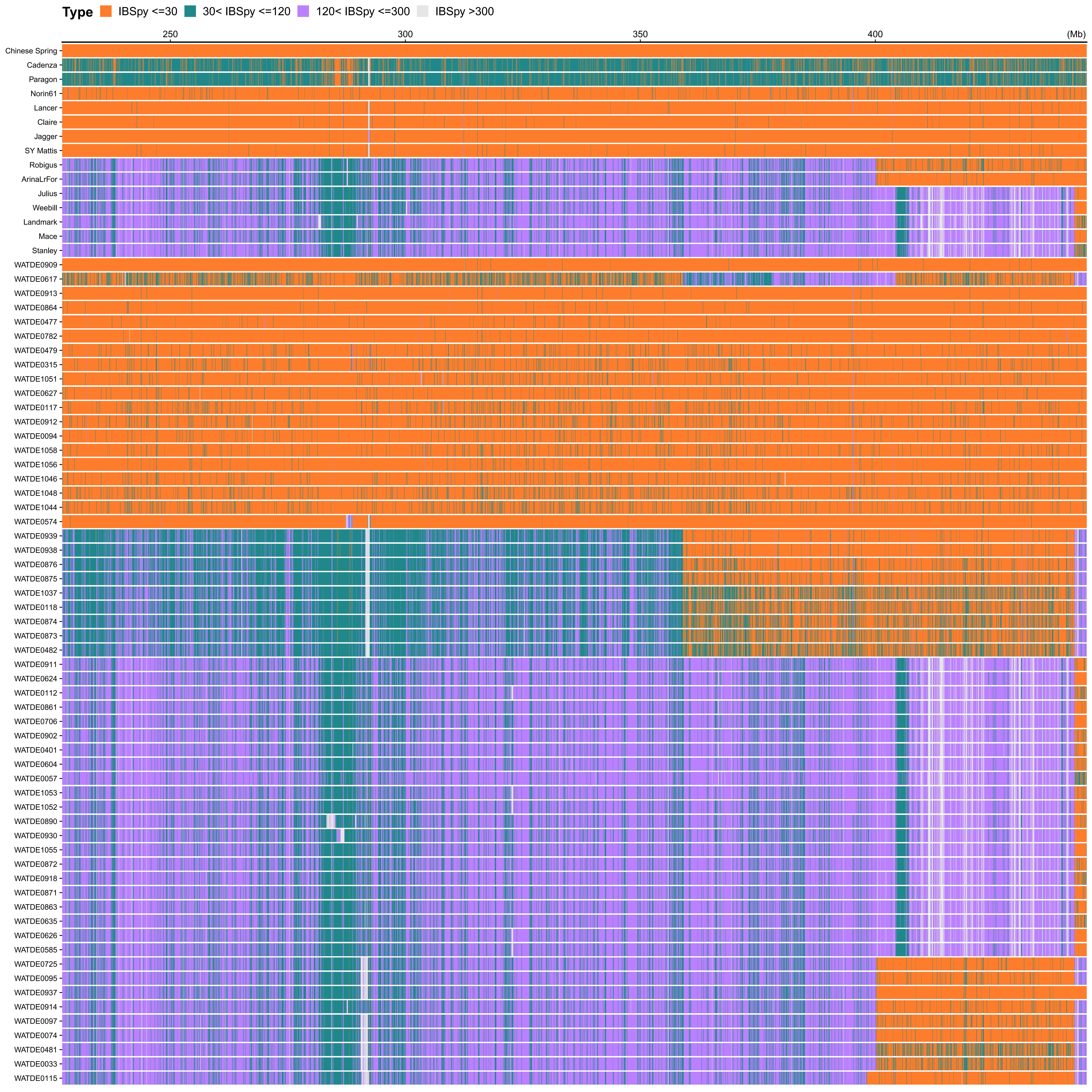

### Fig. S12.pdf

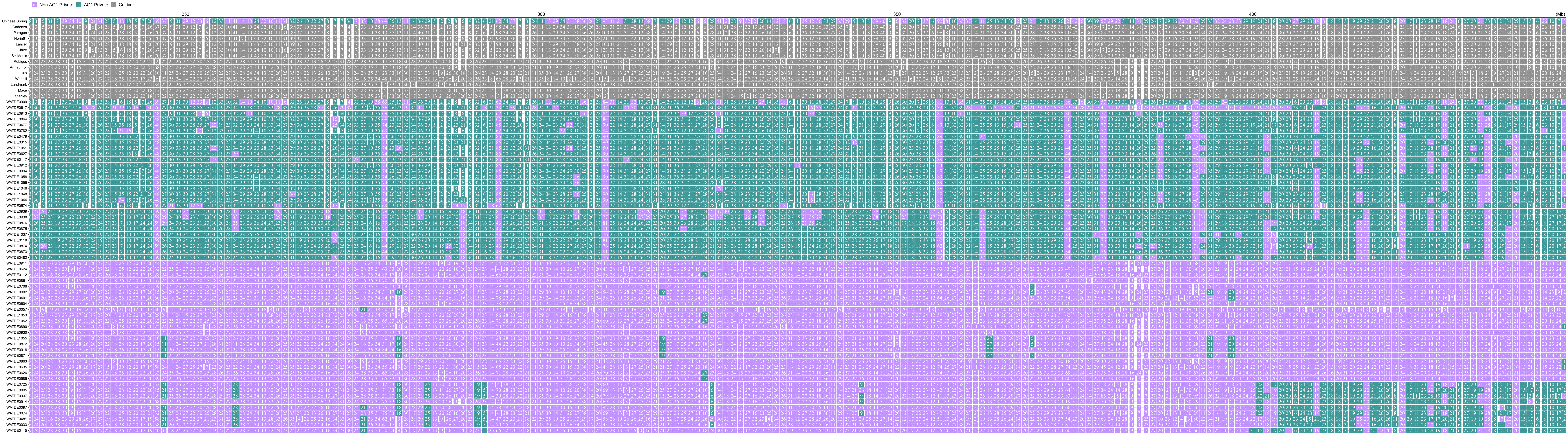

### Fig. S13.pdf

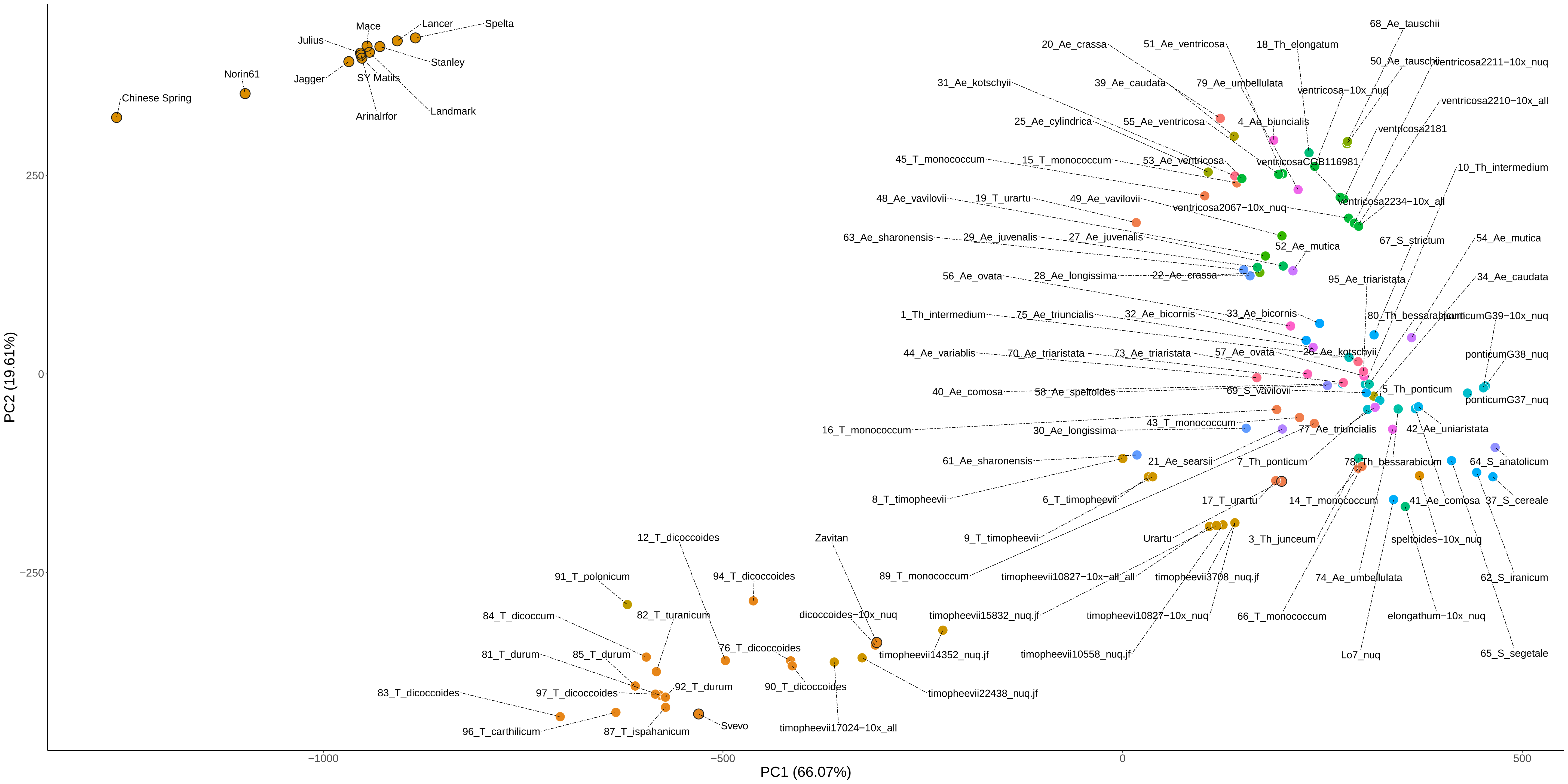

### Fig. S14.pdf

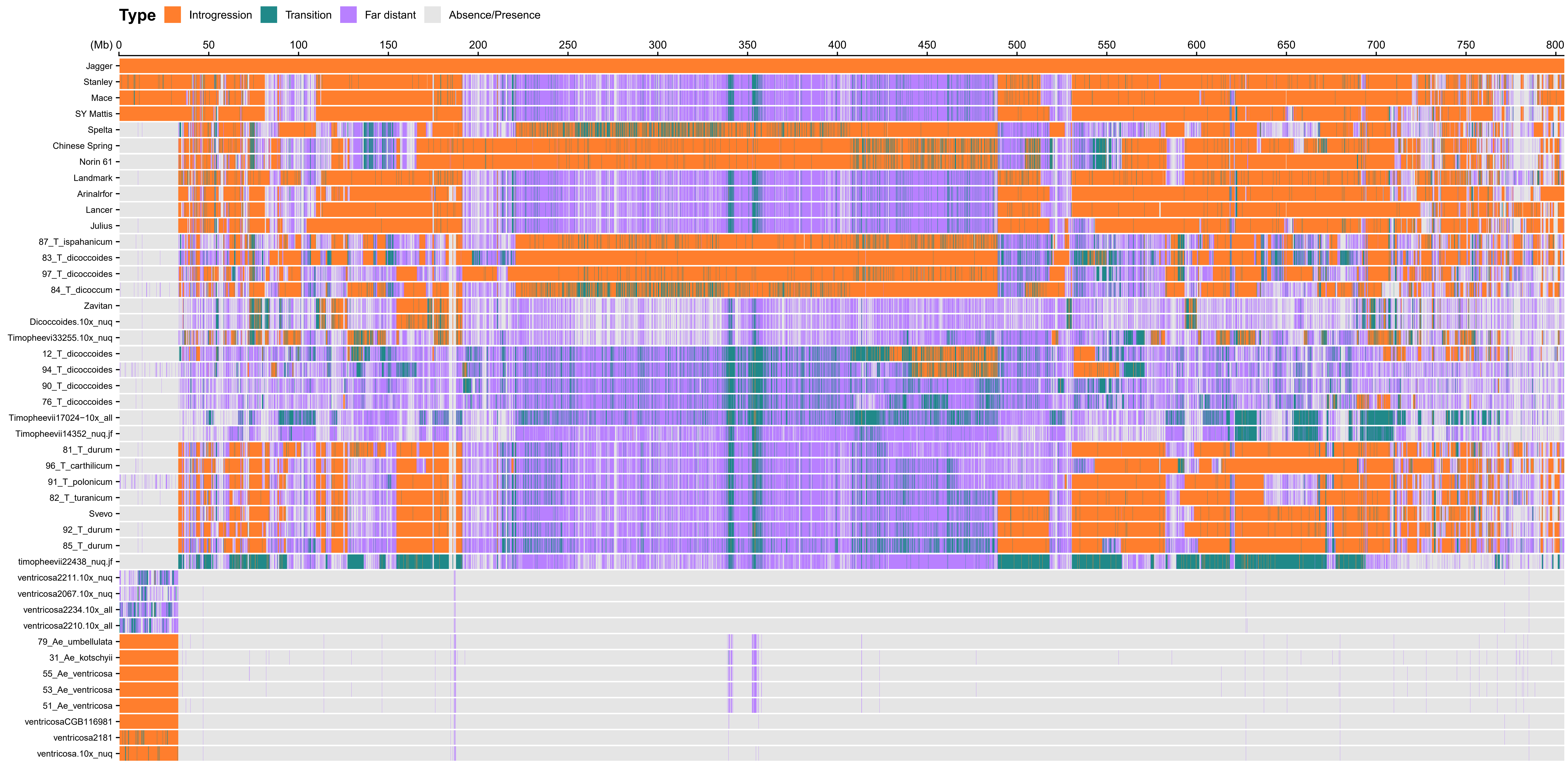

### Fig. S15.pdf

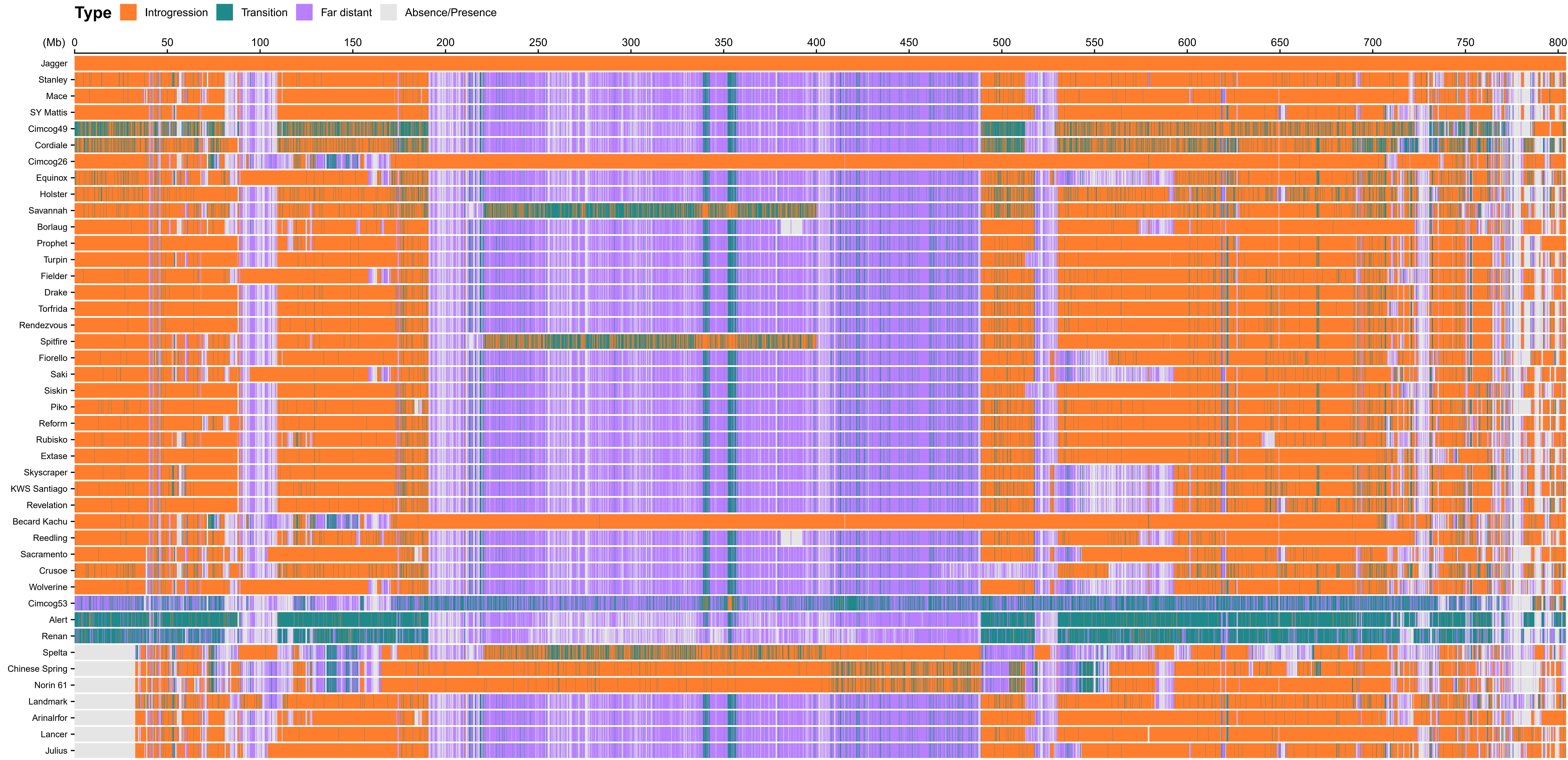

### Fig. S16.pdf

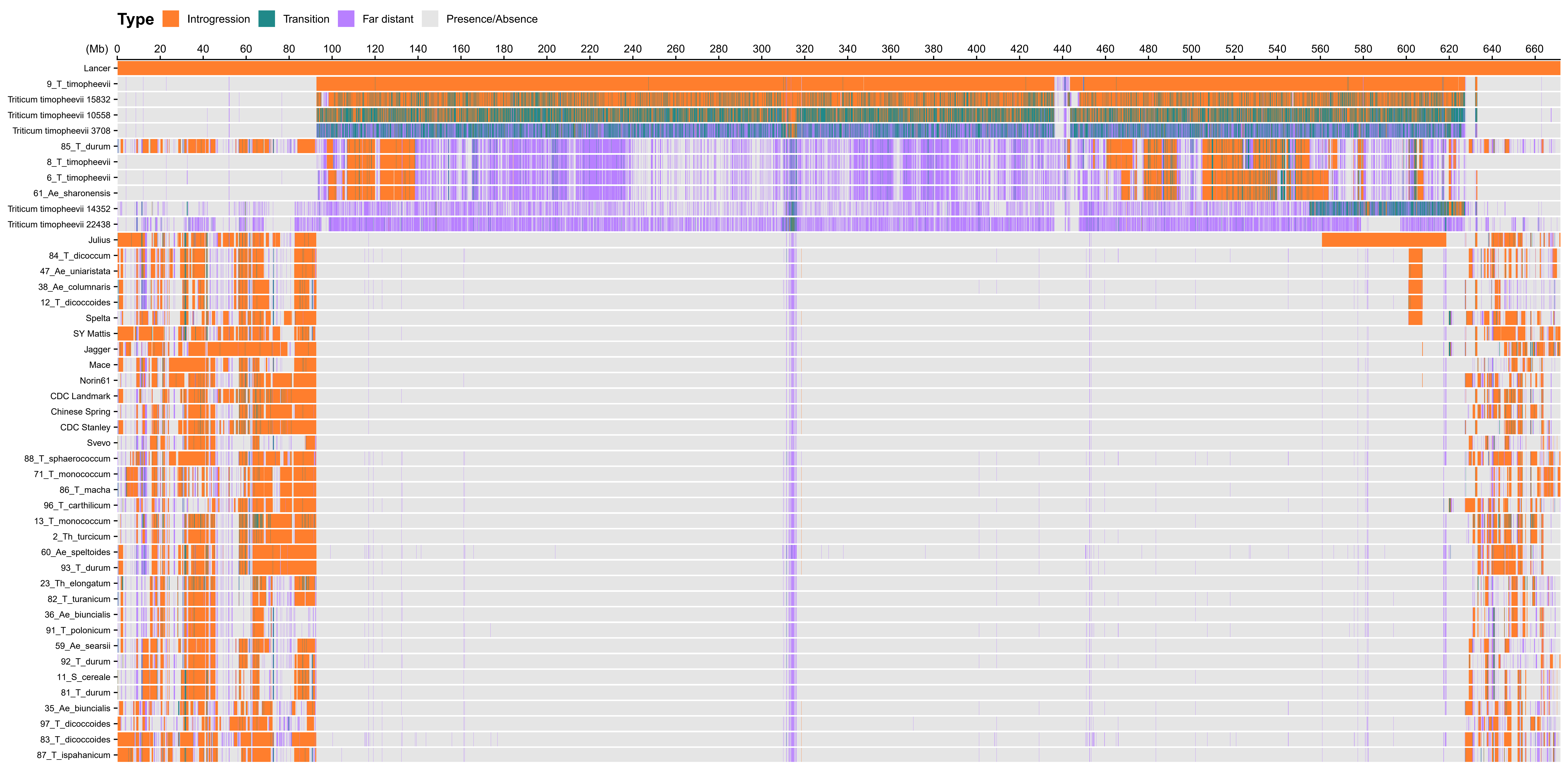

### Fig. S17.pdf

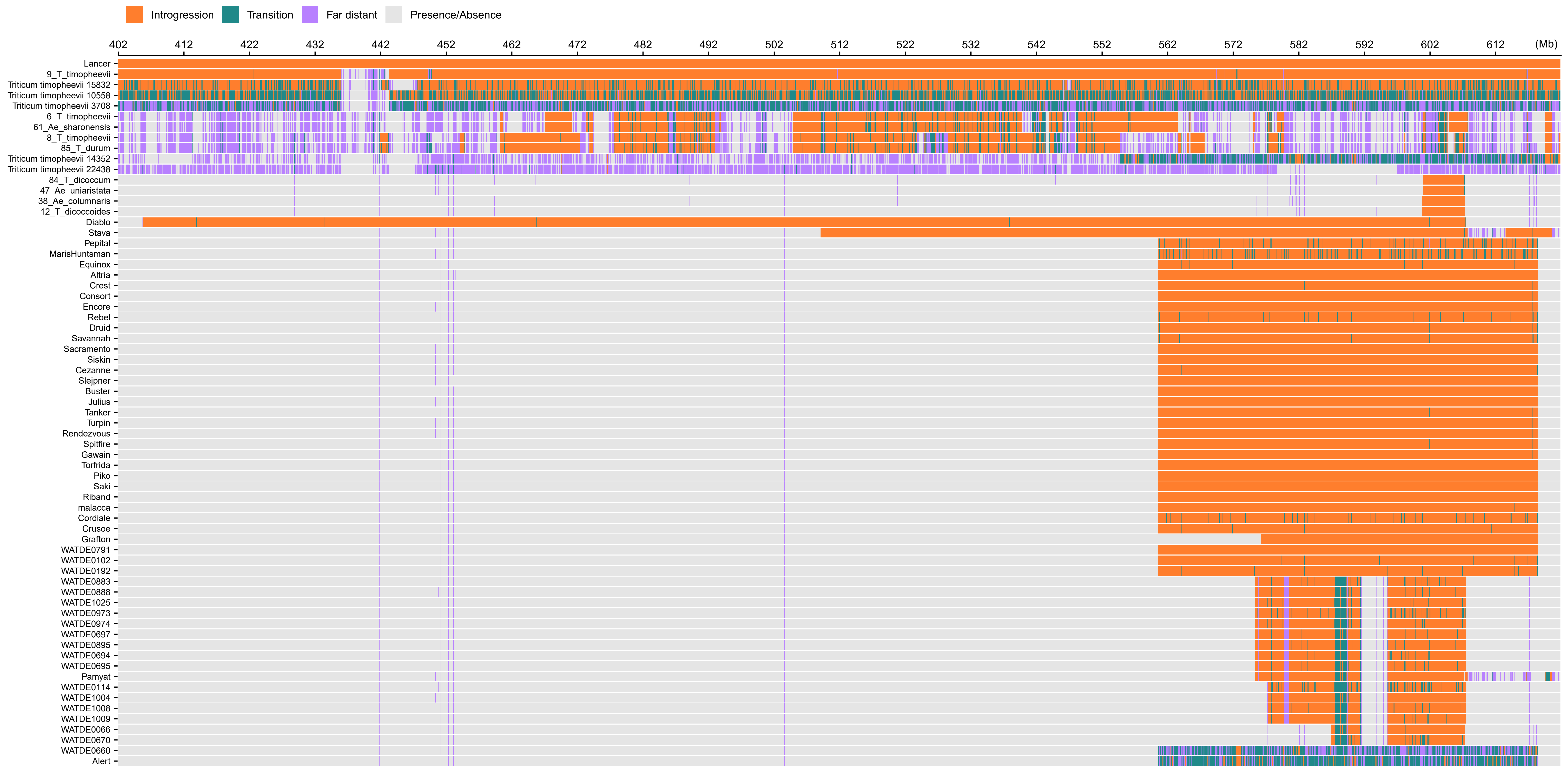

### Fig. S18.pdf

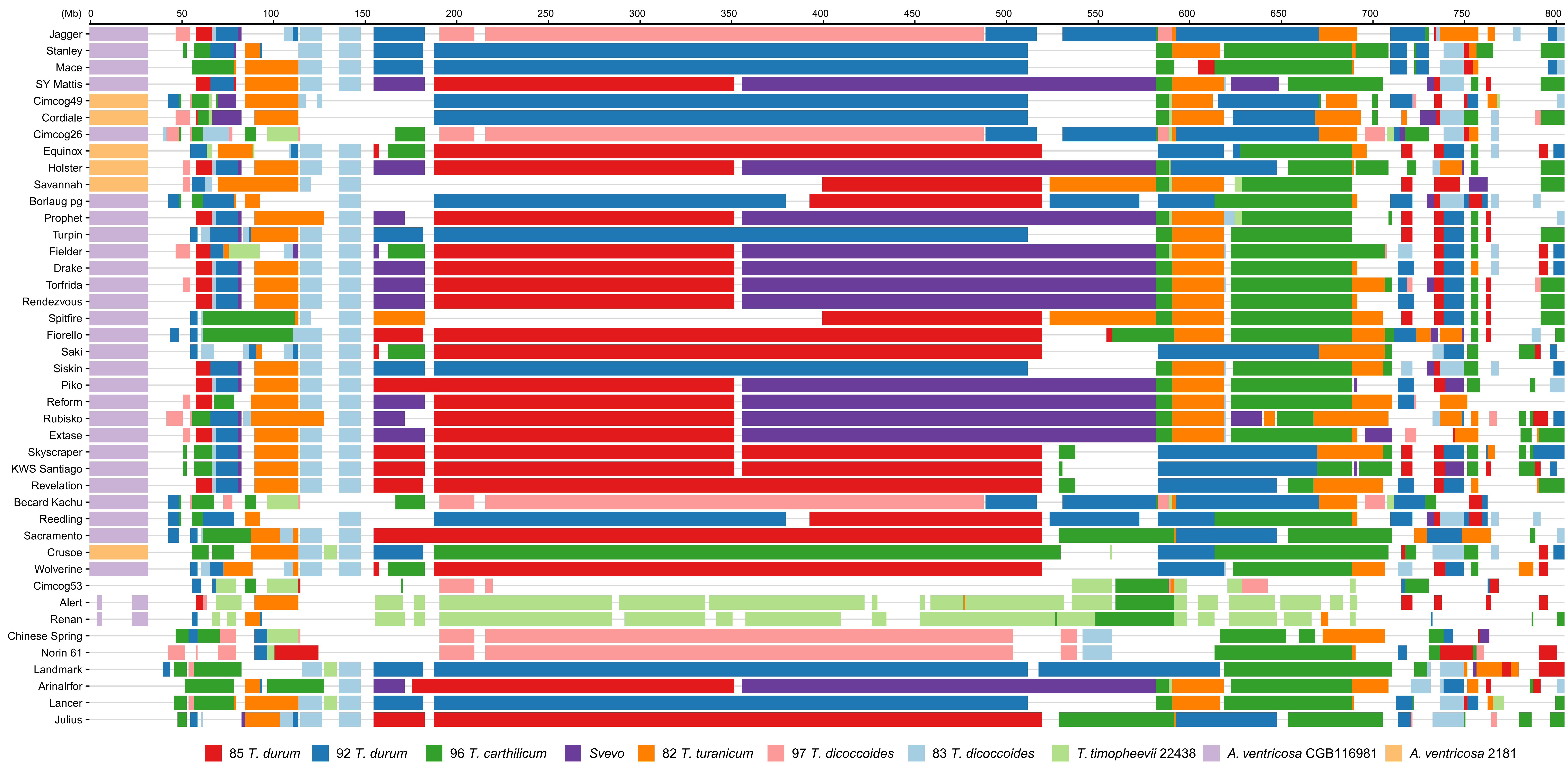

### Fig. S19.pdf

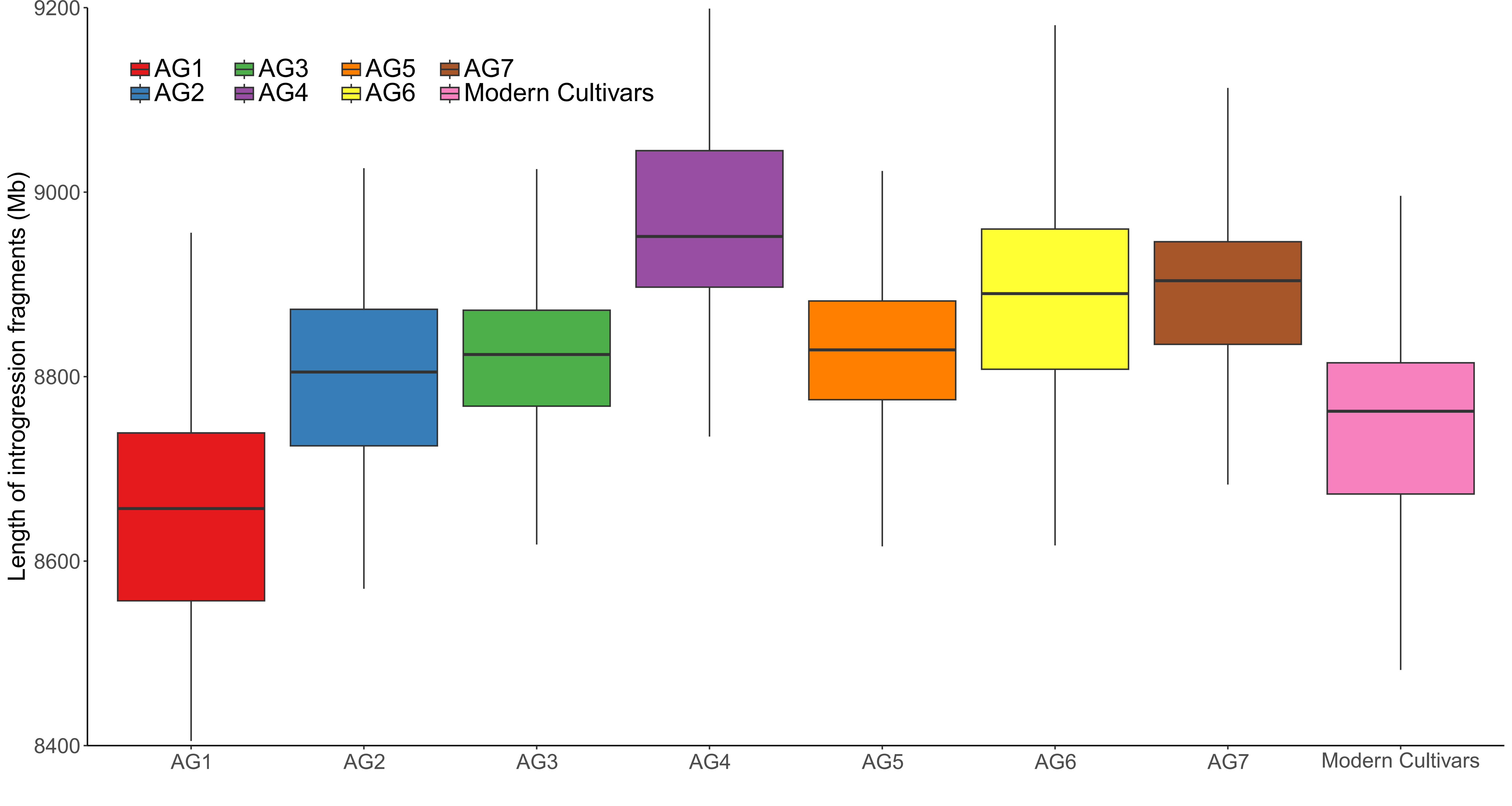

### Fig. S20.pdf

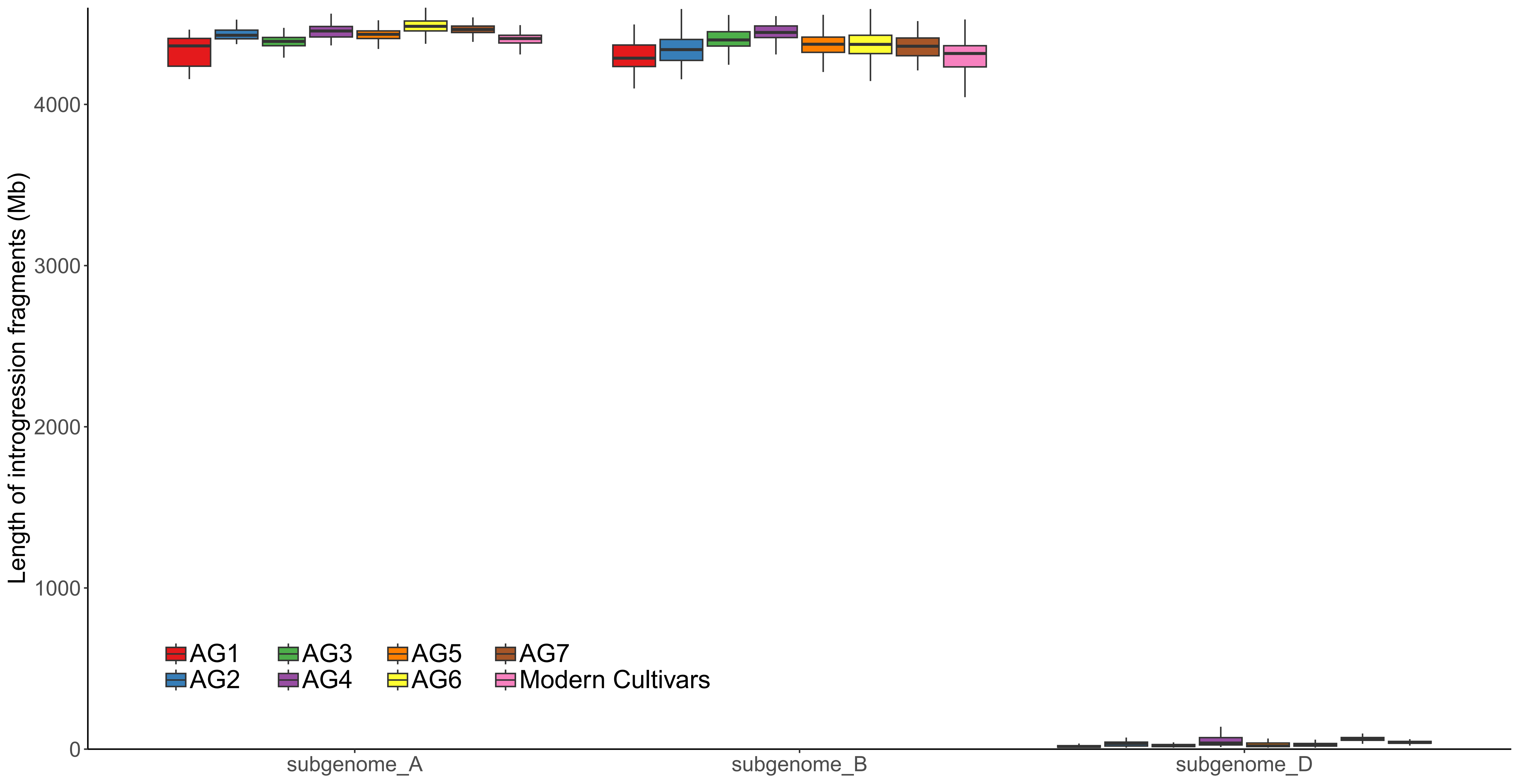

### Fig. S21.pdf

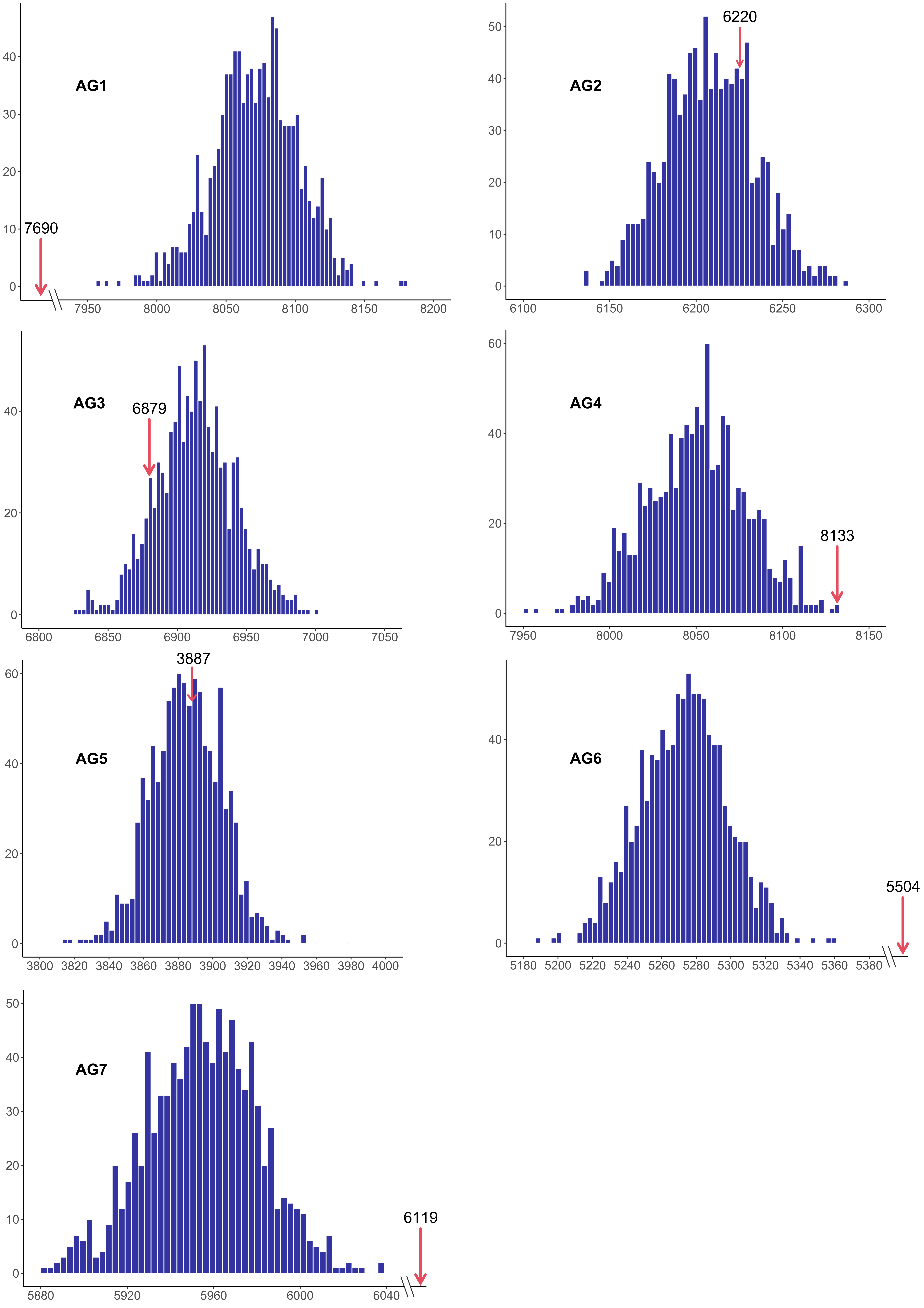

### Fig. S22.pdf

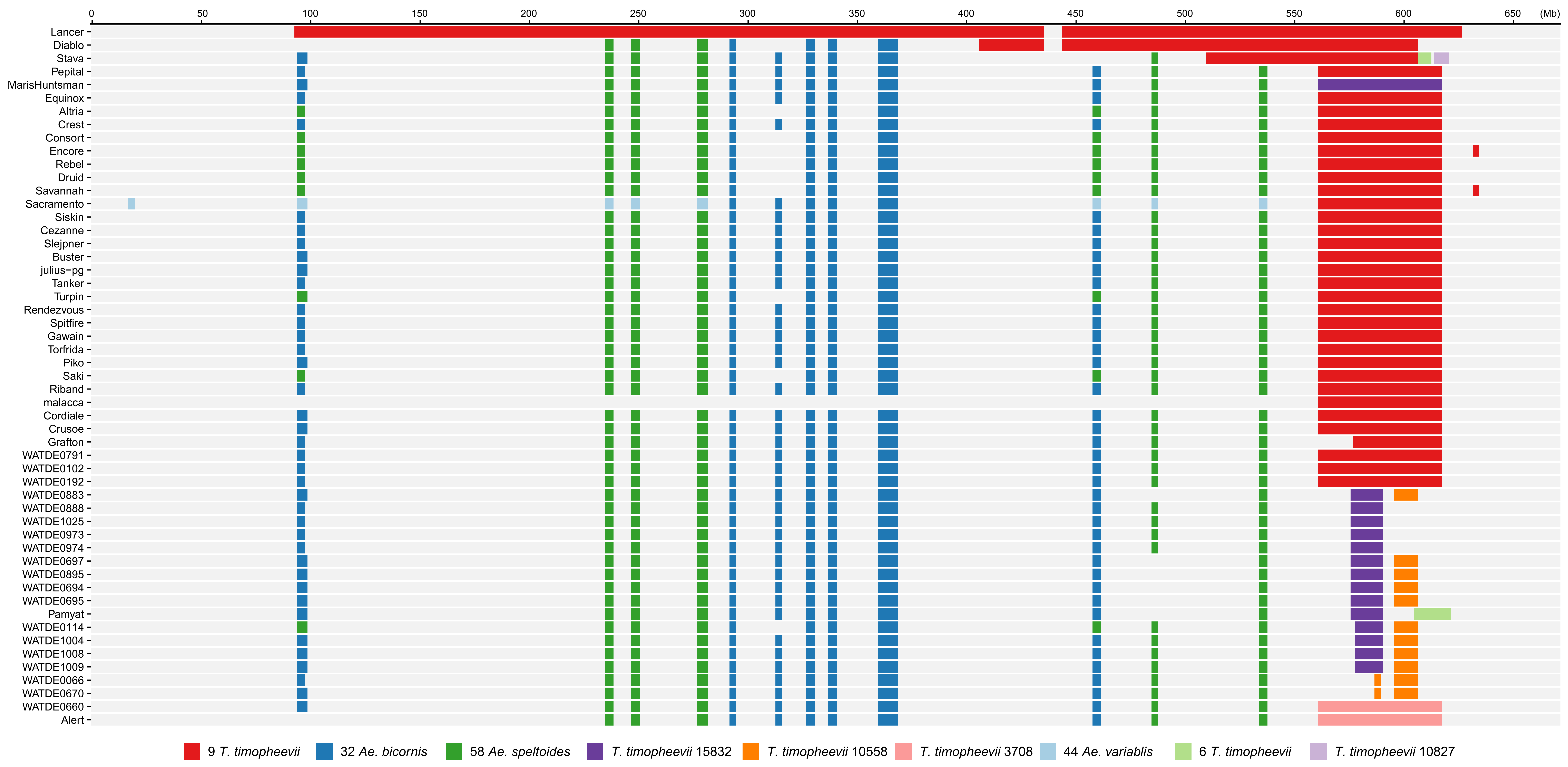

### Fig. S23.pdf

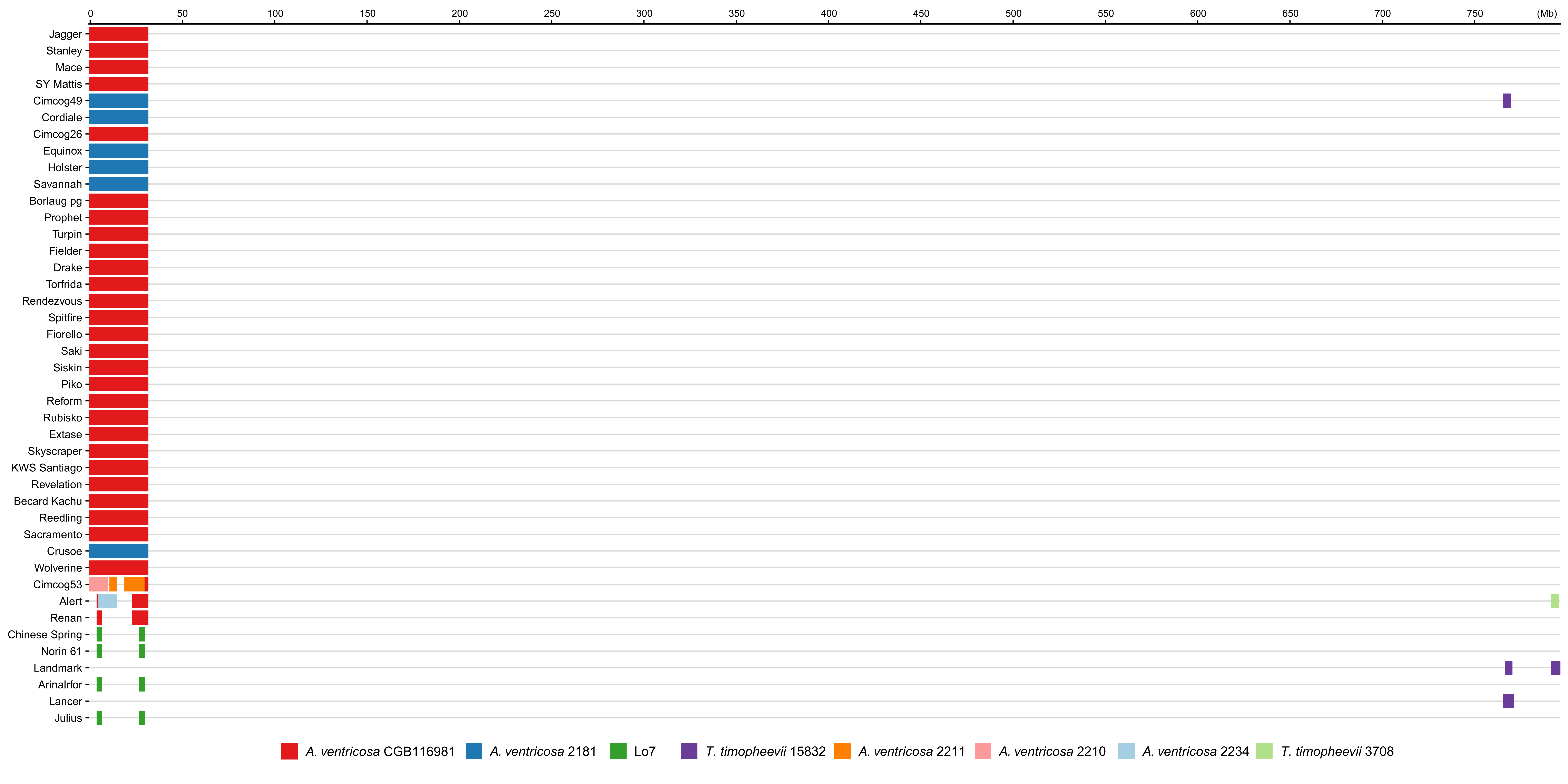

### Fig. S24.pdf

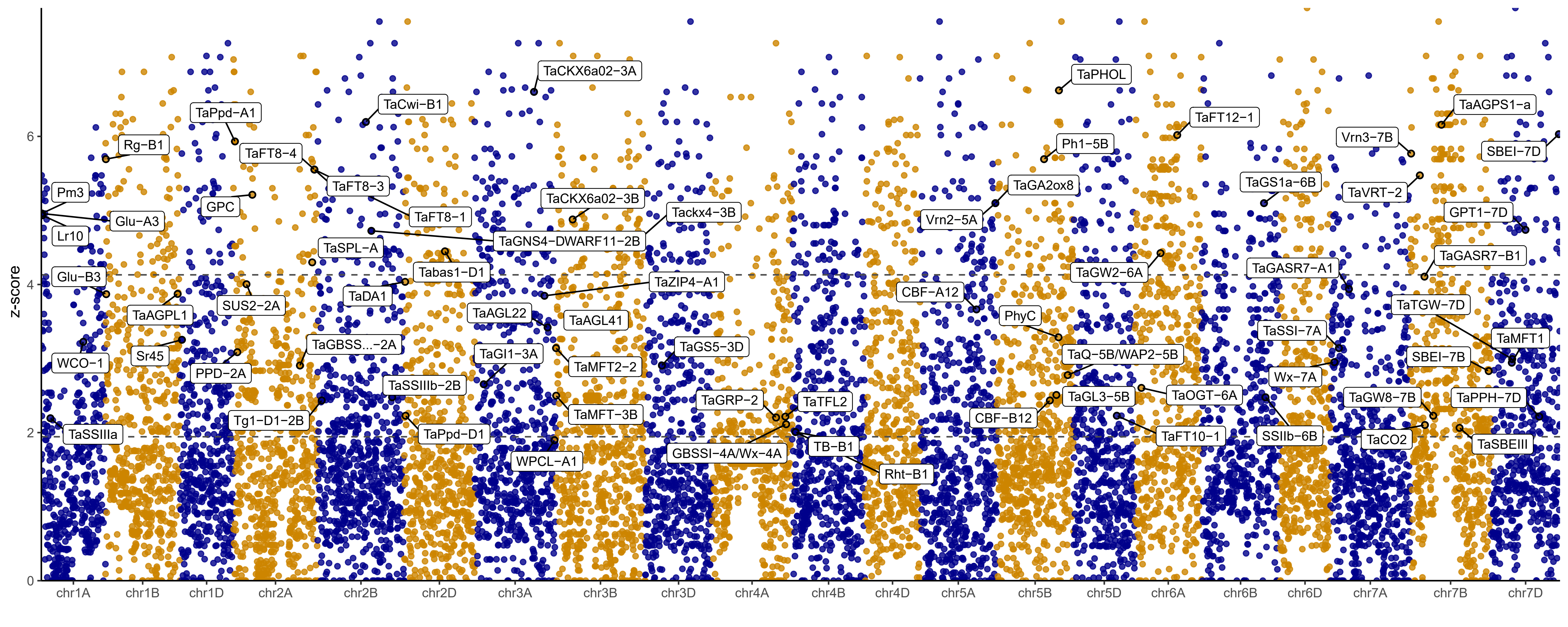

### Fig. S25.pdf

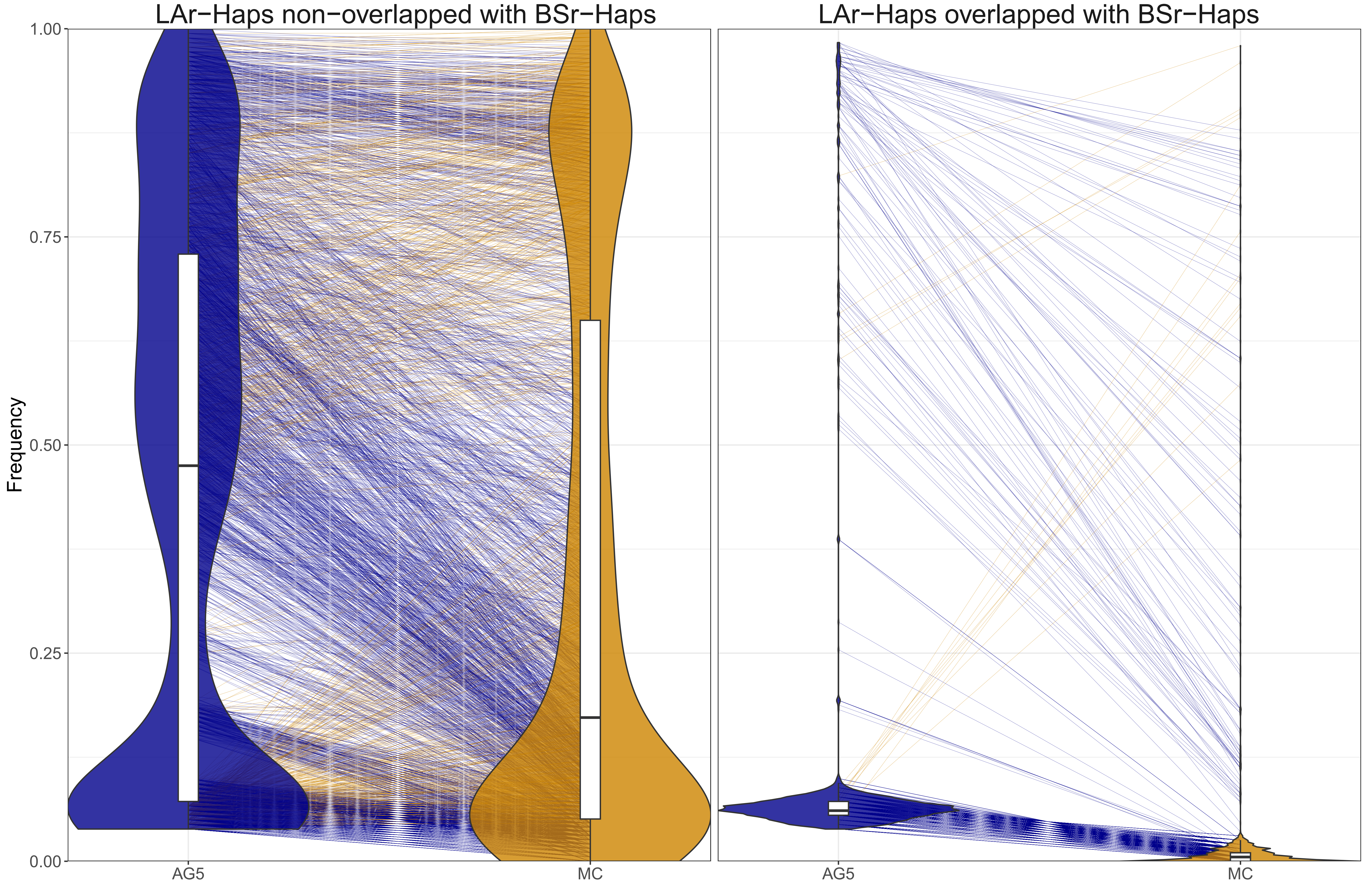

### Fig. S26.pdf

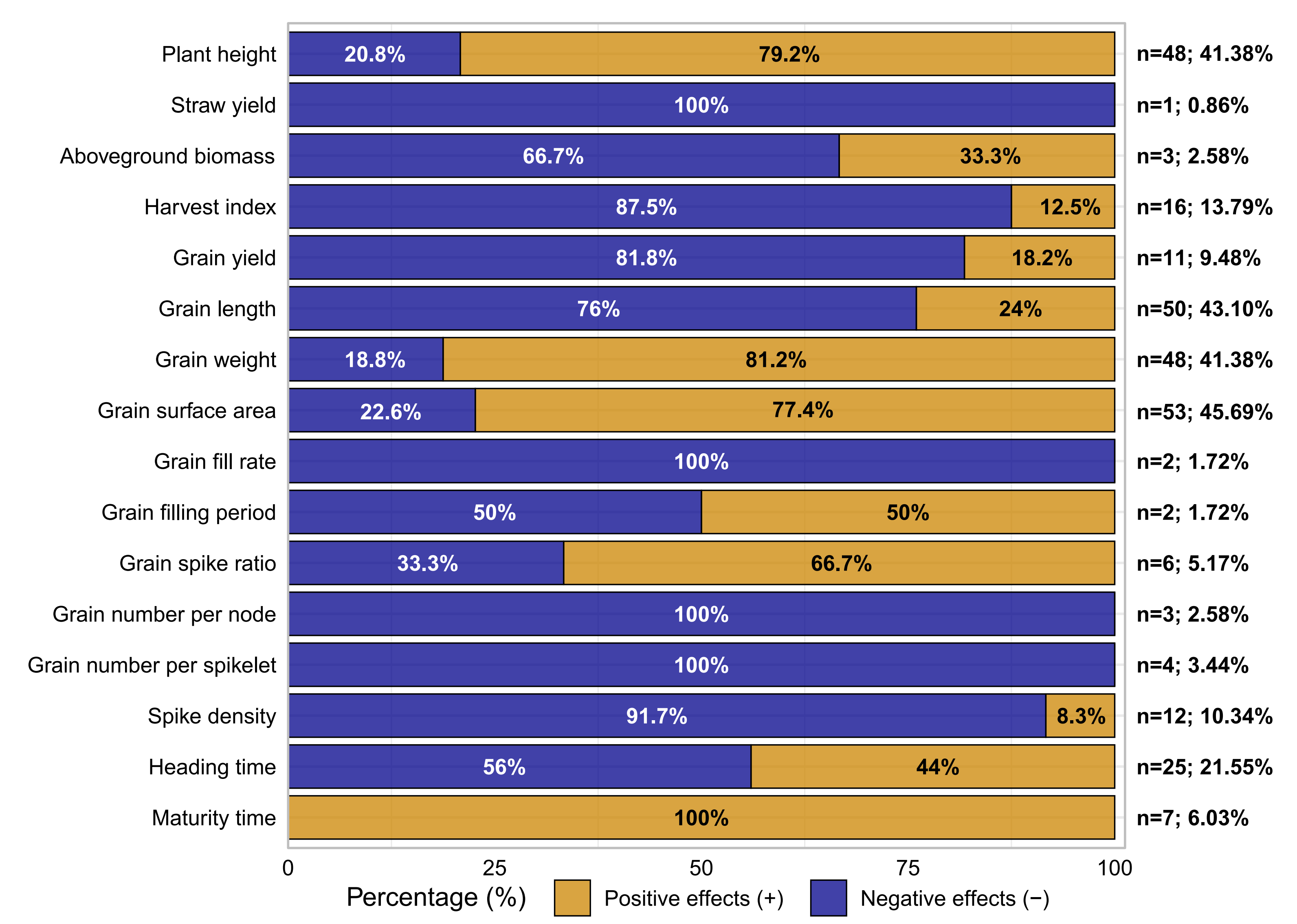
